# Inhibition of Ribosome Biogenesis *in vivo* Causes p53-Dependent Death and p53-Independent Dysfunction

**DOI:** 10.1101/2024.09.25.614959

**Authors:** Charles J. Cho, Thanh Nguyen, Amala K. Rougeau, Yang-Zhe Huang, Sarah To, Xiaobo Lin, Supuni Thalalla Gamage, Jordan L. Meier, Jason C. Mills

**Author notes:** Correspondence: Charles J. Cho, MD, PhD Instructor Section of Gastroenterology & Hepatology Department of Medicine, Baylor College of Medicine One Baylor Plaza, 535EA Houston, Texas 77030, Jason C. Mills, MD, PhD, AGAF Herman Brown Professor of Medicine Co-Director Digestive Disease Center, Chief of Research and Vice Chief, Section of Gastroenterology & Hepatology, Departments of Medicine, Pathology & Immunology, Molecular & Cellular Biology Baylor College of Medicine, One Baylor Plaza, 535E Houston, Texas 77030. Authors contributed equally to this manuscript (Charles J. Cho, Thanh Nguyen). **Disclosures**: The authors declare no conflict of interests.

## Abstract

Ribosomes are critical for cell function; their synthesis (known as ribosome biogenesis; “RiBi”) is complex and energy-intensive. Surprisingly little is known about RiBi in differentiated cells *in vivo* in adult tissue. Here, we generated mice with conditional deletion of *Nat10*, an essential gene for RiBi and translation, to investigate effects of RiBi blockade *in vivo.* We focused on RiBi in a long-lived, ribosome-rich cell population, pancreatic acinar cells, during homeostasis and tumorigenesis. We observed a surprising latency of several weeks between *Nat10* deletion and onset of structural and functional abnormalities and p53-dependent acinar cell death, which was associated with translocation of ribosomal proteins RPL5 and RPL11 into acinar cell nucleoplasm. Indeed, deletion of *Trp53* could rescue acinar cells from apoptotic cell death; however, *Nat10*^Δ*/*Δ^*; Trp53*^Δ*/*Δ^ acinar cells remained morphologically and functionally abnormal. Moreover, the deletion of *Trp53* did not rescue the lethality of inducible, globally deleted *Nat10* in adult mice nor did it rescue embryonic lethality of global *Nat10* deletion, emphasizing p53-independent consequences of RiBi inhibition. Deletion of *Nat10* in acinar cells blocked *Kras*-oncogene-driven pancreatic intraepithelial neoplasia and subsequent pancreatic ductal adenocarcinoma, regardless of *Trp53* mutation status. Together, our results provide initial insights into how cells respond to defects in RiBi and translation *in vivo*.

## INTRODUCTION

Ribosomes are evolutionarily conserved, ubiquitous, essential structures that primarily function in the translation of messenger RNAs (mRNA) into proteins. Investigation into the ribosome biogenesis (RiBi) and mRNA translation processes has been mostly limited to two biological contexts. The first context is *in vitro* studies using rapidly growing (usually tumor-derived) cell lines, where disrupting RiBi or inhibiting translation results in proliferation defects and/or cell death through increased p53 protein levels (Bywater et al., 2012; Farley-Barnes et al., 2019; Mills & Green, 2017; Peltonen et al., 2010; Wei et al., 2018). The second context involves developmental studies, where congenital mutations in specific ribosomal proteins or RiBi components result in organ type-specific defects (e.g., Diamond-Blackfan Anemia or Treacher-Collins syndrome), with various degrees of p53 involvement (Farley-Barnes et al., 2019; Gazda et al., 2012; Jones et al., 2008; Mills & Green, 2017). However, *in vivo* studies on RiBi and mRNA translation processes in adult tissues are severely lacking.

Clinically, RiBi and translation are attractive targets for modulating multiple biological processes, particularly those related to tumorigenesis (Hilton et al., 2022; Khot et al., 2019; Kogure et al., 2013; Nguyen et al., 2023). Disruption of RiBi leads to impaired cell growth and proliferation as well as p53 stabilization, a process dubbed ‘nucleolar stress’ (Boulon et al., 2010). Both RiBi and translation require the direct involvement of hundreds of ribosomal and non-ribosomal proteins, each of which can serve as cell type-specific targets for cancer therapies. To date, we have limited understanding of the ribosome and translation dynamics across different cell types, ranging from constantly proliferating stem cells to fully differentiated cell types (Cho et al., 2024). This gap in knowledge is primarily due to the lack of sufficient tools and modalities for comprehensively testing each RiBi component *in vivo*.

Here, we developed a mouse model to help study effects of disruption of ribosomal dynamics *in vivo.* We generated mice in which the ATP-dependent acetyltransferase enzyme, N-acetyltransferase 10 (NAT10), can be deleted in a temporally and spatially controlled manner. NAT10 has been shown to be a critical enzyme for RiBi and cellular protein translation. Specifically, NAT10 functions in acetylating two cytidine residues in 18S ribosomal RNA (rRNA), which is critical for proper 18S rRNA biogenesis and 40S/80S ribosome production; and in acetylation of transfer RNAs (tRNAs) that deliver serine and leucine to the translating ribosome (Satoshi Ito et al., 2014; S. Ito et al., 2014; Sas-Chen et al., 2020; Sharma et al., 2015). Using this model in combination with mouse models of tumorigenesis with genetic lineage tracing, we comprehensively evaluated how acinar cells in the pancreas, and in mice as a whole, respond to the loss of NAT10 in various developmental and disease-specific contexts.

We observed a surprising latency between the loss of NAT10 and onset of any structural and functional rearrangement in acinar cells. We also observed eventual cytosol-to-nucleoplasm translocation of RPL5 and RPL11, two well-known ribosomal proteins that reportedly stabilize p53 (Castillo Duque de Estrada et al., 2023; Donati et al., 2013; Sloan et al., 2013), in the acinar cells *in vivo*, correlating with p53-induced cell death. Furthermore, p53 deletion rescued *Nat10*-deficient acinar cell death; however, these *Tp53*^Δ*/*Δ^*; Nat10*^Δ*/*Δ^ acinar cells were still deficient in amylase production, revealing that RiBi regulation of cell survival can be uncoupled from its requirement for physiological cell function (e.g., generation of digestive enzymes). Finally, mice with acinar cell-specific expression of mutant, constitutively active *Kras* showed substantially less pancreatic intraepithelial neoplasia (PanIN) and subsequent pancreatic ductal adenocarcinoma (PDAC) when *Nat10* was also deleted, even in the absence of p53.

## RESULTS

### Targeting ribosomal DNA transcription to evaluate ribosome biogenesis and translation *in vitro* and *in vivo*

To establish an optimal system for inhibiting RiBi *in vivo*, we first examined ways to inhibit RiBi *in vitro* using a gastrointestinal cancer-derived cell line, AGS. We tested widely used RNA-polymerase I (pol-I) inhibitor, BMH-21, which suppress ribosomal DNA (rDNA) transcription. BMH-21 treatment (1 μM) resulted in a significant decrease in the nascent 45S pre-rRNA level (Fig. 1A, left) within three hours, while the mature 18S rRNA level remained constant (Fig. 1A, right), consistent with previous reports (Guner et al., 2017; Jacobs et al., 2022). Immunofluorescent (IF) staining showed a reduction in signal of nucleolar proteins involved in multiple steps of RiBi, including UBTF, a marker for the nucleolar fibrillar center (Fig. 1B, left), and NPM1 (Fig. 1B, middle), a marker for the granular component of the nucleolus, as well as their dislocation from the nucleolus to form a nucleolar cap (Fig. 1B, right) (Colis et al., 2014). Decreased cell viability, as measured by the alamarBlue Assay, was noted within 24 hours (Fig. 1C). Similar effects were observed in the colon cancer derived cell line, LS-174T, where a topoisomerase II (TOP2) inhibitor, CX-5461 (100 nM), which can indirectly inhibit pol-I (Bruno et al., 2020; Pan et al., 2021; Xu & Hurley, 2022), caused similar dislocation of the nucleolar protein, TCOF1, and formation of the nucleolar cap (Fig. 1D), along with decreased cell viability (Fig. 1E).

**Fig. 1.**
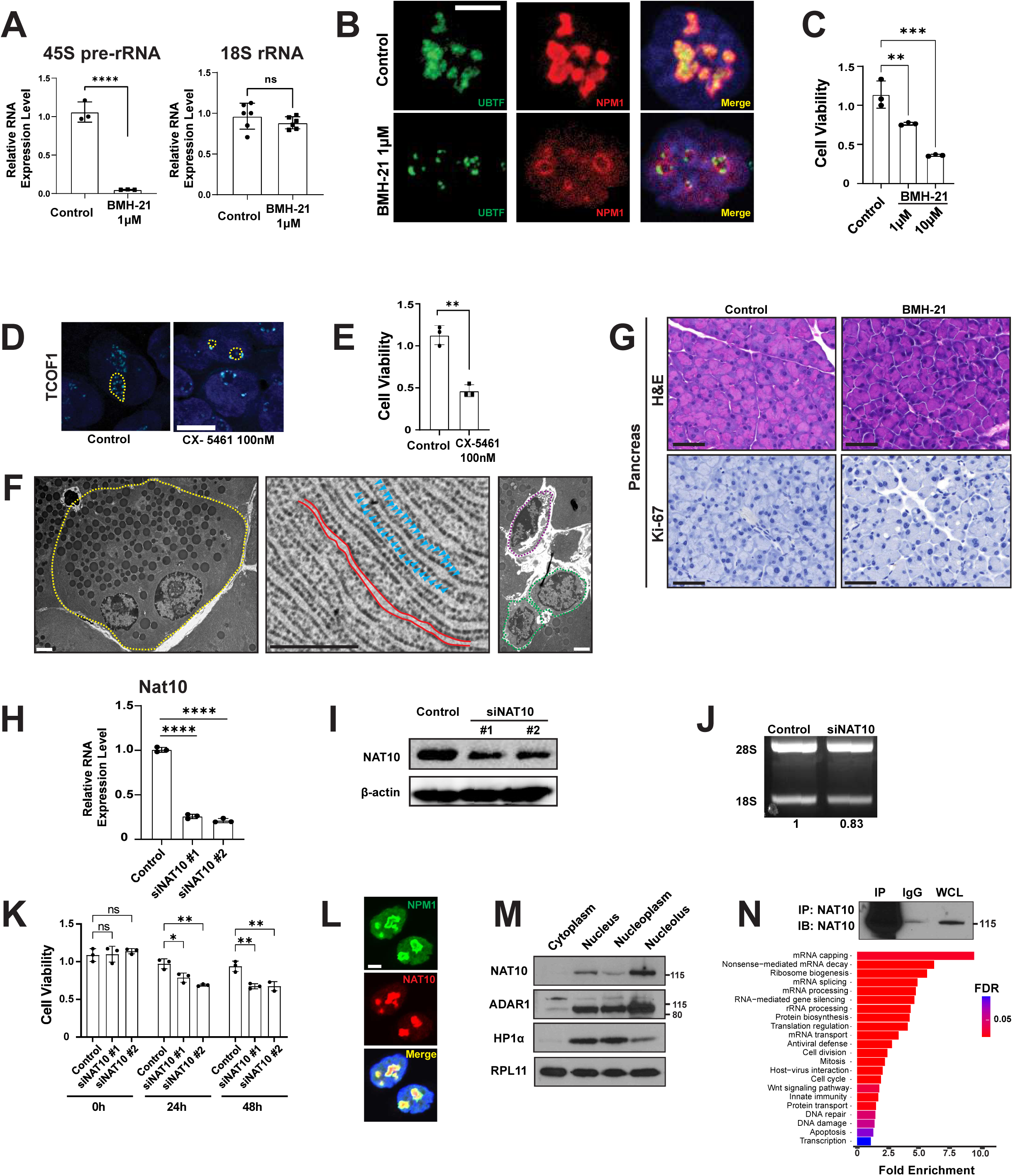
Determination of optimal modalities to study ribosome biogenesis (RiBi) and translational deficiencies in in vitro and in vivo. (A-E) A decrease in 45S pre-ribosomal RNA (rRNA, A, left), a reduction in signal and dislocation of nucleolar proteins UBTF and NPM1 (B), and decreased cell viability (C) are noted in AGS cells treated with BMH-21, a bona fide RNA polymerase I inhibitor. Similar findings are noted in the LS-174T cell line with CX-5461 by immunofluorescence (IF) (D) and cell viability assays (E). (F-G) Comparison of the size of pancreatic acinar cell (yellow, left panel) with endothelial cells (purple, right panel) and ductal cells (green, right panel), scale bar: 2 µm. Rough ER membrane (red, middle panel) and ribosomes (blue arrowheads, middle panel) highlight the abundance of ribosomes in pancreatic acinar cells, scale bar: 500 nm. No obvious changes were noted in routine H&E staining or Ki-67 immunohistochemistry staining of mouse acinar cells in the pancreas upon BMH-21 treatment *in vivo* (G); scale bar: 50 µm. (H-K) Inhibition of NAT10 effectively inhibits RiBi *in vitro*. Knockdown of NAT10 with two different siRNAs in the AGS cell line results in a decrease in NAT10 level as determined by qRT-PCR (H) and western blot (I), which results in a decrease in the 18S/28S rRNA ratio (J) and a decrease in cell viability (K). (L-N) NAT10 is a nucleolar protein involved in RiBi. IF shows NAT10 localizes to the nucleolus and nucleoplasm (L, scale bar: 5 µm), and in subcellular fractionation conducted in the AGS cell line (M). Immunoprecipitation (IP) of NAT10-interacting partner proteins using an anti-NAT10 antibody shows enrichment in pathways related to RiBi and translation over nonspecific IgG pulldown (N); FDR: False Discovery Rate.

Next, we investigated whether targeting rDNA transcription could help assess effects of RiBi inhibition in pancreatic acinar cells *in vivo*. We chose to focus on acinar cells because they are rarely mitotic and long-lived cells that are devoted to their physiological function of translating proteins on abundant rough endoplasmic reticulum (ER), storing those proteins (mostly digestive enzymes) in secretory granules, and secreting them in a regulated manner when they are needed for digestion (Fig. 1F, left and middle). Thus, like cancer cell lines in tissue culture, they have a need for constant translation; however, unlike tumor cells, they largely don’t divide and are governed by finely-tuned physiological function. We again used BMH-21, because it has been shown at the doses we used to inhibit tumor xenograft growth in mice (Peltonen et al., 2014). Using varied BMH-21 dosing schemes based on the previous report (once daily for 5 days vs. twice daily for 3 days, 50 mg/kg, intraperitoneal injection), we did not observe significant morphological changes of pancreatic acinar cells (Fig. 1G, top). We also did not observe any changes in proliferation index, with acinar cells retaining their usual quiescence (Fig. 1G, bottom). Although the lack of phenotype seen *in vivo* with this drug might be explained by insufficient drug delivery or other technical issues, it could also be that the reason for the dramatic effects *in vitro* is because cells in tissue culture (or in xenografts of cancer cells) may need constant RiBi to feed cell growth in order to enter the cell cycle, whereas acinar cells are already large cells and are mitotically quiescent so they have an enormous supply of extant ribosomes on rough (ER) (Fig. 1F-G) (Marcel et al., 2015). The large ribosome pool may ‘buffer’ the effects of blocking RiBi for an extended period of time. To understand impact of loss of RiBi on long-lived cells in an organ, we reasoned we would need a way to stably block RiBi for extended periods *in vivo* in a cell-type-specific way (e.g., via a genetically engineered mouse model).

### NAT10 as a tool to evaluate ribosome biogenesis and translation *in vitro* and *in vivo*

To block RiBi stably and genetically, we focused on NAT10, an RNA acetyltransferase that acetylates two cytidine residues in 18S rRNA as well as serine and leucine transfer RNAs (tRNAs). We hypothesized that deleting the *Nat10* gene would disrupt RiBi by inducing defective 18S rRNA biogenesis and impairing 40S/80S ribosome production, a process that has been well-validated *in vitro* (Satoshi Ito et al., 2014; S. Ito et al., 2014; Sas-Chen et al., 2020; Sharma et al., 2015). We first validated the effectiveness of targeting NAT10 for perturbing RiBi *in vitro*, again using AGS cells. siRNA against NAT10 reduced *Nat10* mRNA (Fig. 1H) and protein as expected (Fig. 1I). Also as expected, loss of NAT10 correlated with reduced RiBi, as evidenced by decreased 18S rRNA levels (Fig. 1J) and decreased cell viability (Fig. 1K). The NAT10 protein was primarily localized in the nucleolus by both IF (Fig. 1L) and subcellular fractionation (Fig. 1M), consistent with the distribution of its known substrates (rRNA and tRNA). Further, immunoprecipitation of NAT10 using an anti-NAT10 antibody followed by mass spectrometry revealed an enrichment in proteins encoding RiBi and translation functions (Fig. 1N). Together, these data confirmed the role of NAT10 as a critical protein involved in the proper maturation of ribosomes and protein translation *in vitro*.

*In vivo*, NAT10 was ubiquitously expressed in the nucleolus and nucleoplasm of all secretory cells examined, including chief cells in the stomach corpus, acinar cells in the pancreas and salivary glands, as well as in other differentiated cell types across various organs such as in the proximal and distal convoluted tubules of the kidney (Fig. 2A). We next confirmed that RiBi is active in pancreatic acinar cells at homeostasis using the 5-Ethynyl uridine (EU) labeling assay, which incorporates into nascent RNAs. Given that the majority of RNAs in a cell are rRNAs, an intense signal in the nucleolus was observed within 2 hours after EU injection (Fig. 2B). In acinar cells, NAT10 was located in the nucleolus (overlapping with the nucleolar protein, NPM1) and in the nucleoplasm (Fig. 2C). Furthermore, NAT10 interacted with the 40S small subunit ribosomal component, RPS6, in the pancreas (Fig. 2D). In sum, these data indicate the role of NAT10 in RiBi and translation *in vivo* aligns with its function observed *in vitro*.

**Fig. 2.**
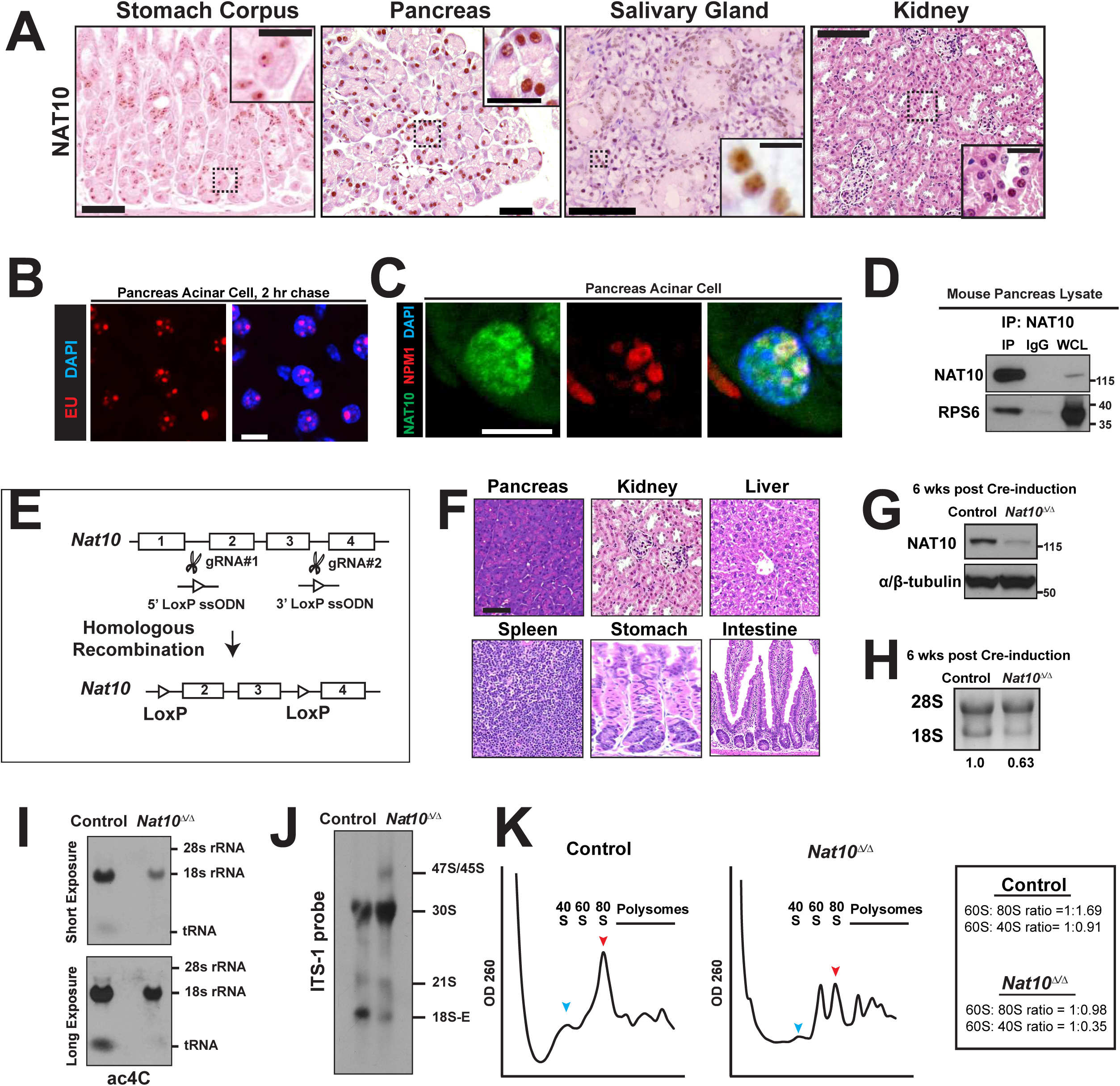
Generation and validation of *Nat10* conditional knockout mouse. (A-D) NAT10 is ubiquitously expressed *in vivo*. The expression of NAT10 was noted in multiple secretory cell types, including chief cells in the stomach corpus (A, far left; scale bars: 50 µm, inset 20 µm), acinar cells in the pancreas (A, left; scale bars: 100 µm, inset 50 µm) and the salivary gland (A, right; scale bars: 50 µm, inset 10 µm), as well as other organs such as in the kidney (A, far right; scale bars: 100 µm, inset 20 µm). The stomach corpus was stained with eosin only for better visualization. 5-Ethynyl uridine (EU) labeling shows nucleolar staining on IF, demonstrating active RiBi in the pancreatic acinar cells (B; scale bar: 10 µm). Co-IF of NAT10 in pancreatic acinar cells show NAT10 overlapping with NPM1, another nucleolar protein, as well as its expression in the nucleoplasm (C; scale bar: 10 µm). IP assays using pancreas lysates demonstrate the interaction of NAT10 with RPS6, a component of the small subunit of the ribosome (D). (E-K) Generation and functional validation of a novel *Nat10* conditional knockout mouse. Two loxP sites were inserted in introns 1 and 3 through homologous recombination, using CRISPR-Cas9 technology (E). The *Nat10^flox/flox^* mice did not show any discernible phenotype across any of the examined organs (F). Cre induction in *Nat10^flox/flox^*; *CAGG-Cre^ERTM^* mice resulted in a sustained decrease in NAT10 protein levels (G), as well as reductions in 18S rRNA levels and the 18S/28S rRNA ratio (H). A decrease in ac4C in 18S rRNA and tRNA (I) and a defect in the biogenesis of mature 18S rRNA (J) was accompanied by a decrease in the amount of the 40S small subunit (blue arrowhead) and 80S assembled ribosomes (red arrowhead) while the 60S large subunit, which do not require 18S rRNA for its formation, remained unaffected (K).

### Generation and validation of *Nat10* conditional knockout mice for assessing ribosome biogenesis and translation *in vivo*

We went on to generate a *Nat10* floxed mouse using the CRISPR-Cas9 approach by inserting loxP sites in intron 1 and intron 3 (Fig. 2E). Recombination of the intervening sequence by *Cre* recombinase would delete all known functional *Nat10* isoforms. Thorough examination of these newly generated, floxed-allele-containing mice showed no evidence of unwanted mutation, alternative splicing site generation (via Sanger sequencing, data not shown), or phenotypic abnormality (Fig. 2F). Next, we sought to assess the quantity and quality of rRNAs and ribosomes, as well as their acetylation status when NAT10 was conditionally deleted. For this, we used the kidney, an organ that has characteristics similar to the pancreas (composed mainly of differentiated cells, quiescent at homeostasis, lacks adult stem cells (Humphreys et al., 2011)), but has the advantage of being relatively free from the endogenous RNases that plague analyses of ribosome assembly status in the pancreas. We generated *Nat10^flox/flox^; CAGG-Cre^ERTM^* mice, where the CAGG-Cre allele is a “universal driver” designed to allow tamoxifen-inducible recombination in most cell types in most adult organs, including those in the kidney (Hayashi & McMahon, 2002). After tamoxifen, we observed an efficient reduction in NAT10 levels (Fig. 2G), a decrease in 18S rRNA levels while 28S rRNA levels remained constant, and a decreased 18S/28S rRNA ratio that persisted at 6 weeks after *Nat10* deletion (Fig. 2H). The presence of N4-acetylcytidine was detected using a specific anti-ac4C antibody (Bryson et al., 2020), which revealed acetylation events occurring in the 18S rRNA and tRNA bands but not in the 28S rRNA, the level of which substantially decreased with the loss of *Nat10* (Fig. 2I). Corroborating those data, rRNA ac4C sequencing, which boasts the highest sensitivity and specificity for determining the acetylation status of RNA (Thalalla Gamage et al., 2021), also demonstrated lower acetylation of 18S rRNA in *Nat10* knockout mice: 95% and 95% in the two wildtype mice vs. 42% and 64% in *Nat10*^Δ*/*Δ^ mice (data not shown in figures). A northern blot using the ITS-1 probe showed defective processing of 21S rRNA to 18S rRNA during the maturation step (Fig. 2J), consistent with previous *in vitro* data (S. Ito et al., 2014; Sharma et al., 2015). Finally, in sucrose gradient analyses, we confirmed decreased 40S (small subunit) and 80S monosomes, but no change in the 60S peak, which does not use NAT10-regulated 18S. Quantitatively, there was a decrease in the 60S:80S ratio (1:1.69 vs. 1:0.98) and 60S:40S ratio (1:0.91 vs. 1:0.35, Fig. 2K). The data herein demonstrate the utility of the novel NAT10 conditional knockout mice in assessing the phenotypes of adult tissues lacking proper RiBi and translation.

### NAT10-deficient acinar cells undergo cell death correlating with RPL5/RPL11 translocation-associated p53-stabilization

To investigate the effects of disrupting RiBi in acinar cells, we crossed *Nat10^flox/flox^* with *Mist1^CreERT2^* mice (*Mist1* is expressed specifically in long-lived, exocrine secretory cells, in particular acinar cells) (Direnzo et al., 2012; Hess et al., 2011; Shi et al., 2009). As expected, Cre recombination caused NAT10 deletion in pancreas within a week (Fig. 3A). However, despite the rapid decrease in NAT10 expression, the 18S/28S rRNA ratio did not change until five weeks after Cre induction (Fig. 3B). We wondered if the persistence of rRNAs was due to the longer half-life in pancreatic acinar cells than reported rRNA half-life in other organs (e.g., 4 to 5 days in hepatocytes (Loeb et al., 1965; Miller, 1973; Nikolov et al., 1983; Stoykova et al., 1983)). To test this, we performed EU pulse-chase metabolic labeling of RNAs, which allowed us to visualize nascent rRNAs in the nucleolus (2 hr chase; Fig. 3C, far left), followed by their rapid dispersal into the nucleoplasm (5 hr chase; Fig. 3C, left) and eventually into the cytoplasm (16 hr chase; Fig. 3C, middle), accurately reflecting the pattern of rRNA maturation and export. We next compared the cytoplasmic intensity of the EU 9 and 16-day ‘chase’ experiment (Fig. 3C, right and far right) and found no stronger intensity than the cytoplasmic EU intensity at baseline. Therefore, it is unlikely that rRNA may persist up to 4 weeks. We interpret the results to indicate that the majority of 18S rRNA is in the cytosolic, mature ribosomes and that it requires prolonged latency for those to be replaced with the nascent, defective ribosomes generated without NAT10.

**Fig. 3.**
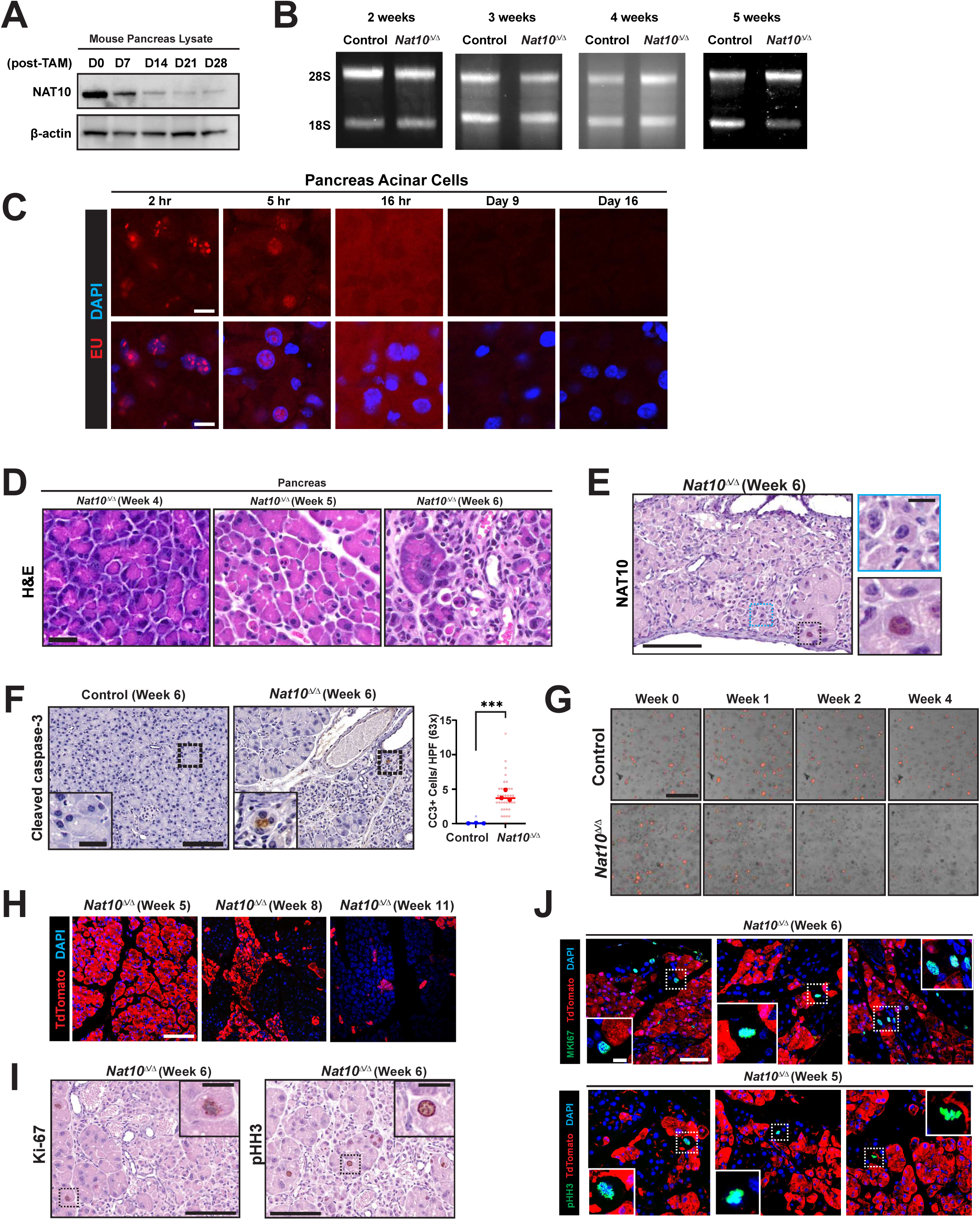
Acinar cells deficient in NAT10 undergo apoptotic cell death after prolonged latency. (A-E) Prolonged latency is noted between loss of NAT10 and induction of the phenotype. Cre induction in *Nat10^flox/flox^; Mist1^CreERT2/+^* mice rapidly decreases NAT10 protein expression levels in the pancreas (A), while a decrease in 18S/28S rRNA ratio in *Nat10*^Δ*/*Δ^ is noted only five weeks after Cre induction (B). (C) EU pulse-chase metabolic labeling of RNAs shows signals in the nucleolus (2 hr chase; far left), nucleoplasm (5 hr chase; left), and cytoplasm (16 hr chase; middle), which is largely absent on EU 9- and 16-day chase (right and far right). Pancreatic acinar cells deficient in NAT10 appeared normal on routine H&E staining until 4 weeks post-Cre induction (D, left), lost basophilic staining by week 5 (D, middle), and by week 6, showed shrinkage in volume and occasional cell death (D, right); scale bars: 25 µm. Notably, shrinking cells were exclusively devoid of NAT10 on IHC (E); Scale bars: 100 µm, inset 10 µm. (F-H) Pancreatic acinar cells deficient in NAT10 undergo apoptotic cell death. (F) Shrunken *Nat10*^Δ*/*Δ^ cells are positive for cleaved caspase-3 staining by IHC (F; scale bars: 100 µm, inset 20 µm). (G) Organoids derived from pancreatic acinar cells of control or *Nat10*^Δ*/*Δ^ mice were cultured for up to 4 weeks; scale bar: 1000 µm. (H) Apoptosis of *Nat10*-deficient cells eventually resulted in the loss of their population *in vivo*, as indicated by TdTomato-positivity throughout the 11-week post-Cre induction period; scale bar: 100 µm. (I-J) Compensatory proliferation occurs in non-recombined acinar cells. Cells re-entering the cell cycle were noted in morphologically intact acinar cells (I; scale bars: 100 µm, inset 20 µm). These cells were negative for the TdTomato signal when lineage traced (J; scale bars: 50 µm, inset: 10 µm).

Furthermore, we noticed significant gross or microscopic morphologic differences in pancreatic acinar cells beginning only 5 weeks post-Cre induction. At this point, histopathological analysis showed diminution of the basophilic, hematoxylin staining normally seen in the basolateral acinar cells (Fig. 3D, middle panel) and general disorganization of acinar architecture. Cell volume was decreased, becoming more pronounced the following week (week 6, Fig. 3D, right panel), at which point apoptotic bodies were frequently observed. These changes affected up to 90% of acinar cells, consistent with the recombination rate of *Mist1^CreERT2^*in the pancreas that we and others have observed for other floxed genes and reporter constructs (Direnzo et al., 2012; Hess et al., 2011). The phenotype was restricted to cells lacking NAT10 staining (Fig. 3E, blue box), whereas healthy-looking acinar cells were strongly positive for NAT10 (Fig. 3E, black box), consistent with cell-intrinsic effects of NAT10 deletion.

We confirmed that NAT10-deficient acinar cells underwent widespread apoptotic cell death at 6 weeks after Cre induction, indicated by cleaved caspase-3 positivity (Fig. 3F, 3.96 ± 0.75 vs. 0.02 ± 0.04 apoptotic cells per high-power field, *P* = 0.0008). To further determine if loss of *Nat10* caused cell-autonomous deaths, we established *ex vivo* acinar organoid cultures. Indeed, *Nat10*-deficient organoids (*Nat10^flox/flox^; ROSA26^LSLTdTomato/+^; Mist1^CreERT2/+^*) died after 4 weeks, whereas control acinar organoids (*Nat10^flox/+^; ROSA26^LSLTdTomato/+^; Mist1^CreERT2/+^*) persisted and emitted fluorescence (Fig. 3G) (fluorescence in this case indicating the cells derived from the original *Mist1*-expressing acinar cells). Similarly, *in vivo* lineage tracing showed that the TdTomato-positive *Nat10*^Δ*/*Δ^ acinar population was eventually replaced by regenerative, TdTomato-negative acinar cells that had escaped Cre recombination and *Nat10* deletion (Fig. 3H). In other words, acinar tissue regeneration in *Nat10*^Δ*/*Δ^ pancreas (Fig. 3I) mainly occurred through compensatory proliferation in adjacent, non-recombined acinar cells, which were distinguished by lack of TdTomato signal (Fig. 3J).

We next interrogated the mechanism underlying pancreatic acinar cell death caused by RiBi inhibition. The aberrant regulation of RiBi or translation can result in the stabilization of p53 through MDM2 sequestration, also known as the ‘nucleolar stress’ theory (Deisenroth & Zhang, 2010; James et al., 2014). However, whether and how different cell types *in vivo* relay specific injury signals to stabilize p53 is largely unknown, and the study of how RiBi, p53, and cell death are related in long-lived cells *in vivo* is almost wholly unknown. Congenital mutations of ribosomal or RiBi-related proteins (known as ribosomopathies) result in p53-dependent developmental defects in only a few selected organs, suggesting the situation *in vivo* is far more complex than the usually rapid p53-induced cell death that results from blocking RiBi in cultured cells. Thus, it will likely be necessary to evaluate the p53 phenotype in a cell-type and tissue-specific context (Farley-Barnes et al., 2019; Mills & Green, 2017). In the pancreas, we determined that p53 was barely detectable in non-injured acinar cells (Fig. 4A and 4B, first lane). Up to four weeks after Cre induction, p53 was unchanged by western blot (Fig. 4B, right four lanes). However, by 5 to 6 weeks after loss of NAT10, p53 expression increased by western blot (Fig. 4C), and p53-positive cells could be seen exclusively within the decreasing *Nat10*^Δ*/*Δ^ acinar population, whereas NAT10-positive wildtype cells were p53-negative (Fig. 4D and 4E). Perhaps the most important regulator of p53 activity in cultured cells is MDM2 which governs p53 degradation and has both active (∼90kDa) and cleaved forms (∼60kDa), where the cleaved MDM2 forms after full-length MDM2 contacts and ubiquitinates p53 (Huschtscha et al., 2009; Pochampally et al., 1999). In non-injured pancreas, both forms were detectable by western blot (Fig. 4F, first lane). Up to four weeks after Cre induction, MDM2 were unchanged by western blot (Fig. 4F, right four lanes). By 5 to 6 weeks after loss of NAT10, we did not notice an alteration in the total amount of full-length MDM2 on western blot (Fig. 4G); however, the cleaved form of MDM2 decreased in pancreas lysates from *Nat10*^Δ*/*Δ^ mice, supporting the interpretation that there is loss of p53-MDM2 interaction. In addition, occasional acinar cells at 6 weeks that were positive for MDM2 on IHC were exclusively nucleoplasmic (Fig. 4H). Together, the data suggest changes in the pattern of MDM2-p53 interaction starting at 5 weeks after RiBi blockade.

**Fig. 4.**
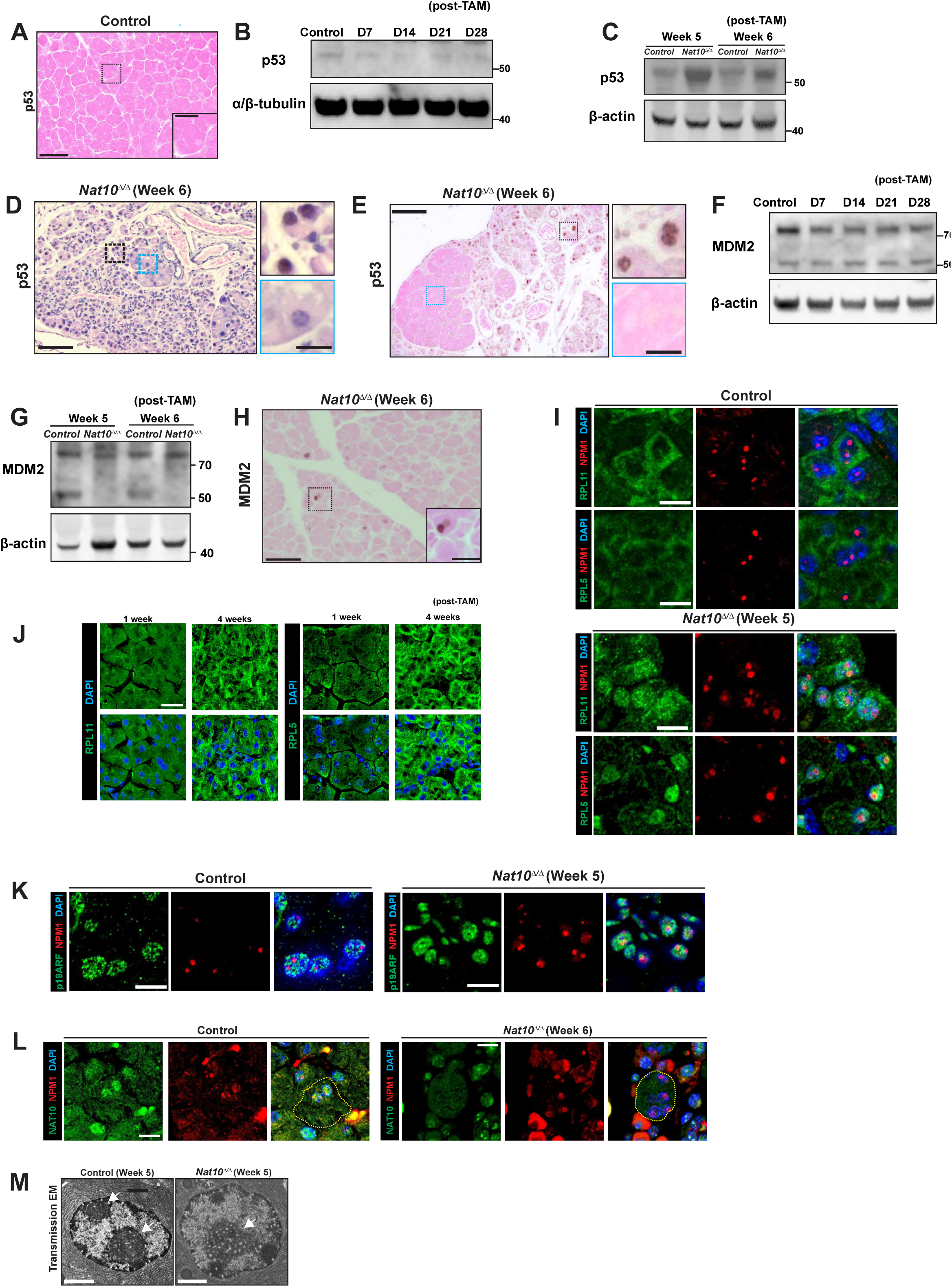
Apoptotic cell death in *Nat10*-deficient acinar cells correlates with p53 stabilization that is associated with RPL5 and RPL11 translocation. (A-E) Non-injured pancreatic acinar cells lack p53, as shown by IHC (A; scale bars: 50 μm, inset 20 μm, stained with eosin only for better visualization) and western blot (B, far left lane), the similar finding of which is noted in *Nat10*^Δ*/*Δ^ mice up to 4 weeks after Cre recombination (B, right four lanes). An increase in p53 level is noted, corresponding to the time when acinar cells undergo microscopic morphologic changes (5-6 weeks), as shown by western blot (C) and IHC (D&E, E image with eosin only for better visualization; scale bars: 50 μm, inset 10 μm). (F-H) Both intact and cleaved forms of MDM2 are expressed in non-injured pancreas (F; far left lane), the pattern of which is relatively unaltered up to 4 weeks after Cre recombination (F, right four lanes). At 5-6 weeks after Cre induction, a decrease in the amount of cleaved MDM2 is noted, while the intact MDM2 amount is relatively unchanged (G). A nuclear MDM2 staining pattern is noted (H, scale bars: 50 μm, inset 20 μm) on IHC. (I-K) RPL5 and RPL11, components of the 5S ribonucleoprotein (RNP) complex, mainly localize to the cytoplasm in pancreatic acinar cells in control mice (I, top; scale bar: 10 μm) and *Nat10*^Δ*/*Δ^ acinar cells up to 4 weeks post-Cre induction (J, scale bar: 20 μm) In contrast, *Nat10*^Δ*/*Δ^ acinar cells show enrichment of RPL5 and RPL11 in the nucleoplasm, 5 weeks after Cre induction (I, bottom; scale bar: 10 μm). p19ARF consistently shows a nucleoplasmic pattern in both groups (K, scale bar: 10 μm). (L-M) Nucleolar size and function are retained in *Nat10*^Δ*/*Δ^ acinar cells. IF of NPM1 shows localization to the nucleolus in *Nat10*^Δ*/*Δ^ acinar cells, similar to controls (L; scale bar: 10 μm). Transmission electron microscopy (EM) show the nucleolus (arrows) in *Nat10*^Δ*/*Δ^ acinar cells with a ‘honeycombing’ appearance (M; scale bar: 2 µm) without obvious reduction in nucleolar size.

The nucleoplasmic translocation of various ribosomal and nucleolar proteins including RPL5, RPL11, NPM1 and p14ARF/p19ARF has been reported during RiBi disruption (Castillo Duque de Estrada et al., 2023; Donati et al., 2013; Kurki et al., 2004; Lindström et al., 2000; Sloan et al., 2013; Zindy et al., 1998). Hence, we examined the subcellular localization of RPL5 and RPL11. In the pancreas of the control mice, both ribosomal proteins were enriched in the cytoplasm (Fig. 4I, top) as was noted in *Nat10*^Δ*/*Δ^ mice of up to 4 weeks after NAT10 deletion, (Fig. 4J). In contrast, by 5 weeks, there was a dramatic shift of RPL5 and RPL11 to the nucleoplasm with exclusion from the nucleolus and cytoplasm in *Nat10*^Δ*/*Δ^ mice (Fig. 4I, bottom). On the other hand, we did not notice a significant change in the localization of p19ARF (Fig. 4K) and NPM1 (Fig. 4L), which can reportedly mediate nucleolar stress in tissue culture cells (Kurki et al., 2004; Lindström et al., 2000; Zindy et al., 1998). The mechanism operating in acinar cells *in vivo* appears distinct from traditional nucleolar stress such as that induced by pol-I transcription inhibition in cultured cells (Derenzini et al., 1998). Note, for example, *in vitro*, there is functional and morphological disruption of the nucleolus when RiBi is blocked (Fig. 1B and 1D). In *Nat10*^Δ*/*Δ^ acinar cells, the nucleolus does not collapse, and nucleolar proteins such as NPM1 maintain proper localization within the organelle, despite absence of NAT10 (Fig. 4L). Transmission electron microscopy (EM) also demonstrated maintenance of nucleolar structure; however, the development of a multifocal radiolucent area (honeycomb pattern) in nucleoli suggests there is atypical nucleolar sub-compartmental reorganization, which needs to be further characterized (Fig. 4M). Collectively, these data show that pancreatic acinar cells deficient in RiBi and translation eventually undergo p53-dependent cell death. The death correlates with certain (but not necessarily all) aspects of the nucleolar stress pathway defined *in vitro*, namely RPL5/RPL11 relocation to nucleoplasm and changes in MDM2-p53 interactions.

### Loss of *Nat10* first causes ribosomopathy, then causes organellar and secretory dysfunction in pancreatic acinar cells

Because, after long latency, ribosomal proteins dramatically relocalized from cytoplasm to nucleus, and p53-associated cell death eventually occurred after NAT10 deletion, we next asked if a gradual change in the structure and/or function in acinar cells could be detected that might predispose cells to apoptosis. We first noticed decreased abundance of amylase (a key digestive-enzyme cargo protein of acinar secretory granules) at 5 weeks Cre induction (Fig. 5A), specifically in the acinar cells where Cre-recombination had occurred (Fig. 5B). At the same time, we also noticed distortion in the organization of the lamellar rough ER at the ultrastructural level, with decrease in ER stack abundance and with ER luminal widening, patterns consistent with detachment and/or loss of ribosomes. (Fig. 5C, middle panels). By week 8, nearly all *Nat10*^Δ*/*Δ^ pancreatic acinar cells were dead (Fig. 3H), with the few survivors exhibiting near-complete loss of cell volume and ER that had scant ribosomes bound (Fig. 5C, right panels). However, the abundance of these structures was unchanged during the first 4 weeks after RiBi blockade as determined by western blot using: antibodies against the luminal ER protein HSPA5 and amylase for secretory granule cargo (Fig. 5D) and RPS6, RPL5, and RPL11 for ribosomal proteins (Fig. 5E). This, again, contrasted with the early, rapid loss of NAT10 after Cre induction (Fig. 3A), thus further emphasizing the latency between loss of RiBi and effects on cell homeostasis.

**Fig. 5.**
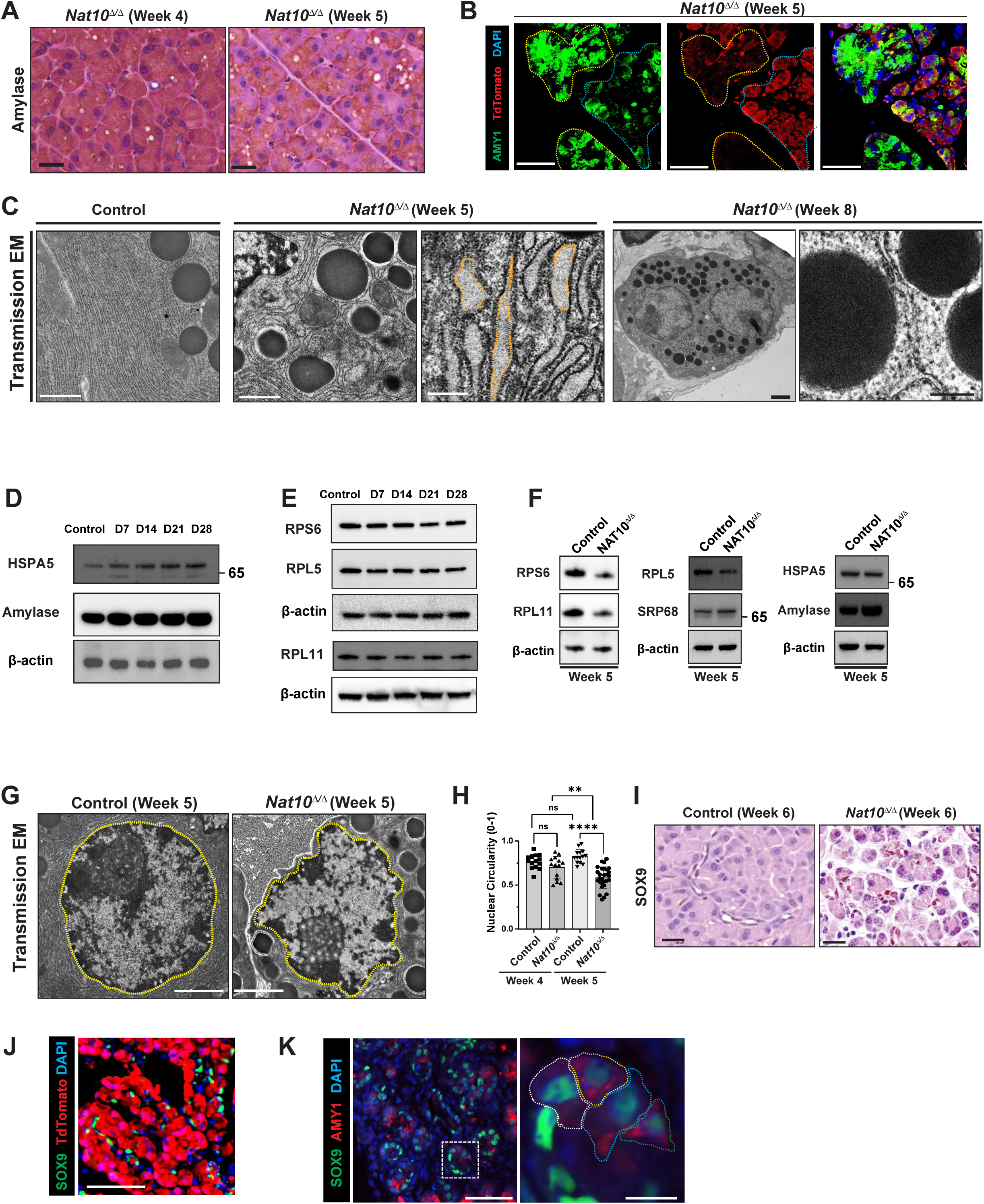
Loss of *Nat10* first causes loss of ribosomes then organellar and secretory dysfunction in pancreatic acinar cells. (A-C) A decrease in amylase, the primary secretory enzyme produced by acinar cells, begins at 5 weeks post Cre induction (A; scale bar: 20 µm), which selectively occurs in acinar cells that underwent Cre recombination and is therefore TdTomato-positive (B, scale bar: 50 µm). Transmission EM images show abundant ribosomes bound to rough ER in the control pancreas (C, left), while *Nat10*-deficinent acinar cells show disorganized, scanty ER with widened ER lumens (yellow lines) 5 weeks after Cre induction (C, middle), with only a few shrunken acinar cells remaining 8 weeks post-Cre induction (C, right). Note that a few ribosomes are still bound to rough ER (C, far right); scale bars: 1 µm, 1 µm, 500 nm, 2 µm, 200 nm, from left to right. (D-F) A decrease in ribosomal proteins but not the non-ribosomal proteins occurs 5 weeks after Cre induction. ER translation-related proteins, HSPA5 and PERK, and amylase (D), as well as ribosomal proteins (E) do not decrease throughout the 4 weeks after Cre induction. A substantial decrease in the amount of both large (RPL5, RPL11, and RPL26) and small (RPS6) subunit ribosomal proteins occurs 5 weeks after Cre induction, while the ratio between ER translation-related proteins and gatekeeper protein remains relatively constant (F). (G-H) The circularity of the nuclear membrane gets disrupted, mimicking a ‘laminopathy’ appearance (G; scale bar: 2 µm). Quantification of nuclear circularity (H). (I-K) *Nat10*^Δ*/*Δ^ acinar cells do not undergo acinar-to-ductal metaplasia (ADM). SOX9 is positive in ductal cells in the pancreas of both control (I, left; scale bar: 20 µm) and *Nat10*^Δ*/*Δ^ mice (I, right; scale bar: 20 µm). SOX9-positive cells in *Nat10*^Δ*/*Δ^ mice are TdTomato-negative, ruling out their acinar cell origin (J; scale bar: 50 µm). Acinar cells undergoing ADM are positive for both amylase and SOX9 (K; scale bars: 50 µm, inset: 20 µm).

If *in vitro* models are faithful proxies for ribosomal behavior *in vivo*, loss of NAT10 should first decrease 40S, then 80S ribosomes, because ribosomal proteins not bound to ribosomes should be quickly degraded through ubiquitination (Sung et al., 2016). Accordingly, after a 5- to 6-week latency post-Cre induction, we did observe decreased abundance of pancreatic ribosomal proteins (Fig. 5F). However, HSPA5, SRP68, and Amylase were relatively unchanged as compared to the housekeeping protein, β-actin, at 5 weeks (Fig. 5F), indicating ribosomal components were primarily and disproportionately affected by loss of NAT10. Finally, at 5-weeks after Cre induction, we noticed an increase in the irregularity of the nuclear membrane (nuclear circularity 0.83 vs. 0.58; *P* < 0.0001, Fig. 5G, 5H for quantification), similar to the laminopathic features seen in NAT10-deficient cells *in vitro* (Sharma et al., 2015).

We next investigated whether *Nat10*^Δ*/*Δ^ acinar cells could undergo acinar-to-ductal metaplasia (ADM). Acinar cells progress to ADM by a stepwise rearrangement of cell architecture known as paligenosis that includes changes in regulation of protein translation and massive rearrangement of cell architecture (Miao et al., 2020; Radyk et al., 2021; Willet et al., 2018). Paligenosis allows acinar cells that escape cell death after tissue to regenerate and replace those cells that died. Induction of SOX9 in acinar cells is a marker of ADM, occurring during the second stage of paligenosis. As expected, in the control pancreas, SOX9 was positive only in normal ductal cells, which are morphologically distinct from those of acinar cells on routine H&E staining (Fig. 5I, left). We also noticed SOX9-positive cell population when NAT10 was deleted (Fig. 5I, right); however, these cells were negative for TdTomato (Fig. 5J), indicative of their ductal origin. In contrast, TdTomato-positive, *Nat10*^Δ*/*Δ^ acinar cells were completely resistant to ADM up to six weeks after Cre induction (Fig. 5J), which contrasts with massive ADM with wildtype acinar cells diffusely inducing SOX9 (Fig. 5K) from a well-characterized model involving injections of the secretagogue cerulein (Brown et al., 2022; Miao et al., 2020; Radyk et al., 2021; Willet et al., 2018). Thus, *Nat10*^Δ*/*Δ^ acinar cells are unable to undergo ADM. The data herein suggest that, although there is surprisingly long latency, the ribosome is the first structure affected by the loss of NAT10 in the pancreatic acinar cells. Effects on multiple other organelles are subsequent.

### Deletion of p53 rescues pancreatic acinar cells from apoptotic death but is unable to rescue loss of cell function

We had noted that acinar cells lacking NAT10 eventually begin to die by apoptosis. They also express p53, have altered MDM2-p53 interactions, and nucleoplasmic accumulation of specific large ribosomal proteins. At about the same time, they exhibit alterations in ultrastructure, size, and function. We next tested if p53 induction only correlated with all the alterations induced by NAT10 loss or was required for them. To investigate this, we generated mice deficient in both NAT10 and p53 in acinar cells (Nat10*^flox/flox^*; Trp53*^flox/flox^*; ROSA26*^LSLTdTomato/+^*; Mist1*^CreERT2/+^*, *Nat10*^Δ^*^/^*^Δ^; *Trp53*^Δ*/*Δ^). We noticed that, relative to *Nat10*^Δ*/*Δ^*; Trp53*^Δ*//+*^ control cells, a significantly higher number of *Nat10*^Δ*/*Δ^*; Trp53*^Δ*/*Δ^ cells re-entered the cell cycle (Ki-67 positive cell number, 10.5/HPF vs. 2.6/HPF, *P* < 0.0001, Fig. 6B, Fig. 6C for quantification) at 6 weeks after Cre induction. There was also a significant increase in the number of cells specifically in G2/M phase (pHH3 positive cell number, 3.7/MPF vs. 0.5/MPF, *P* = 0.0004, Fig. 6D, Fig. 6E for the quantification) as well as significantly less cell death (cleaved caspase-3 positive cell number, 2.1/MPF vs. 0.3/MPF, *P* = 0.02, Fig. 6F, Fig. 6G for quantification). TdTomato-positive acinar cells deficient in both NAT10 and p53 were the principal drivers for this increase in proliferative cells (Fig. 6H, blue box), suggesting loss of p53 altered the NAT10 deletion phenotype. We also noticed that *Nat10*^Δ*/*Δ^*; Trp53*^Δ*/*Δ^ acinar cells had larger nucleoli (Fig. 6I, yellow box, Fig. 6A) versus non-recombined acinar cells (Fig. 6I, blue box), suggestive of active RiBi; however, deletion of p53 did not reverse the decrease in the 18S rRNA level (Fig. 6J). Finally, to determine if loss of p53 in *Nat10*^Δ*/*Δ^ acinar cells also prolonged cell survival, we lineage-traced the fraction of TdTomato-positive acinar cells up to 9 weeks of follow-up. This fraction of recombined cells was consistently higher in the double knockout mice than in NAT10-deficient mice with wildtype p53 (85.1% vs. 26.2%, *P* < 0.0001, Fig. 6K, Fig. 6L for quantification). Thus, p53 is required for the cell death phenotype eventually caused by loss of NAT10. On the other hand, the surviving TdTomato-positive acinar cells in *Nat10*^Δ*/*Δ^*; Trp53*^Δ*/*Δ^ pancreas, largely lacked amylase, indicating that, although they survived, p53- and NAT10-deficient acinar cells could not retain normal physiological function (Fig. 6K). The data show that p53 is a critical mediator of the apoptotic death of pancreatic acinar cells when RiBi is blocked by NAT10; however, p53 is dispensable for the defects in cell architecture and function caused by loss of RiBi.

**Fig. 6.**
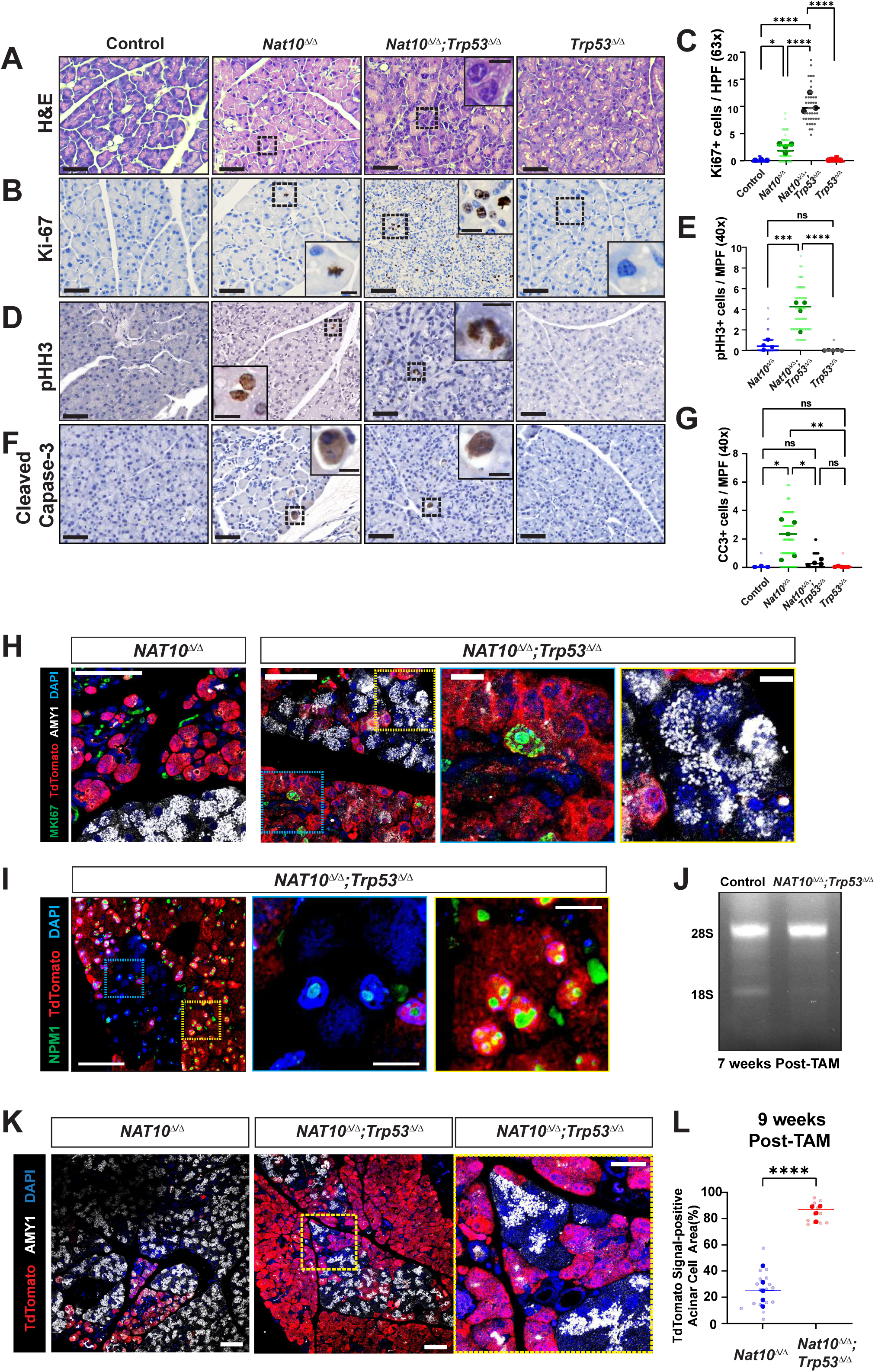
Deletion of p53 rescues pancreatic acinar cells from cell death but not from loss of cell morphology and function. (A-G) Pancreatic acinar cells with conditional knockout of both *Nat10* and *Trp53* (*Nat10*^Δ*/*Δ^*; Trp53*^Δ*/*Δ^) were examined using routine H&E staining (A), proliferative indices (B-E), and apoptosis markers (F-G; scale bars: 50 μm, insets 10 μm). (H-J) Cells in the cell cycle (Ki-67 positive) were mainly *Nat10*^Δ*/*Δ^*; Trp53*^Δ*/*Δ^ cells, as indicated by the positive signal for the TdTomato lineage tracer (H; scale bars: 50 μm, insets 10 μm), where an increase in the nucleolar size was also noted (I; scale bars: 50 μm, insets 10 μm). However, the 18S/28S rRNA ratio in these cells remained decreased (J). (K-L) TdTomato-positive *Nat10*^Δ*/*Δ^*; Trp53*^Δ*/*Δ^ acinar cells not only proliferated but also survived up to 9 weeks post-Cre induction (K), comprising the vast majority of the cell population (L). However, amylase staining was only observed in the non-recombined (TdTomato-negative) acinar cell population (K), suggesting that *Nat10*^Δ*/*Δ^*; Trp53*^Δ*/*Δ^ acinar cells were still defective in amylase production; scale bars: 50 μm (main image), 20 μm (inner box).

### Cell types in different tissues show variable susceptibility to *Nat10* loss, and p53 deletion does not rescue the ribosomopathic phenotype of globally *Nat10*-deficient mice

We next examined the effects of loss of NAT10 other cell types in the murine models to investigate whether loss of NAT10 impacts various cell types to a similar extent. *Mist1^CreERT2^* can induce recombination not only in the pancreatic acinar cells but also in other exocrine secretory cells, including acinar cells in the salivary gland and chief cells in the stomach corpus, which allowed us to evaluate how different cell types respond to *Nat10* loss. Despite efficient Cre recombination in the acinar cells of the submandibular salivary gland (Fig. 7A) and chief cells of the stomach corpus (Fig. 7B), we did not notice consistent alteration in the cell morphology, proliferation, or cell death indices after up to 9 weeks of follow-up in both organs (Fig. 7C and 7D). When these cells also lost *Trp53*, there were similarly no obvious histological, cell cycle or cell death-related changes (Fig. 7E and 7F). Together, these data demonstrate that the susceptibility to inhibition of RiBi *in vivo* may differ among cell types so data from *in vitro* studies cannot be generalized to physiologically functioning cells *in vivo*.

**Fig. 7.**
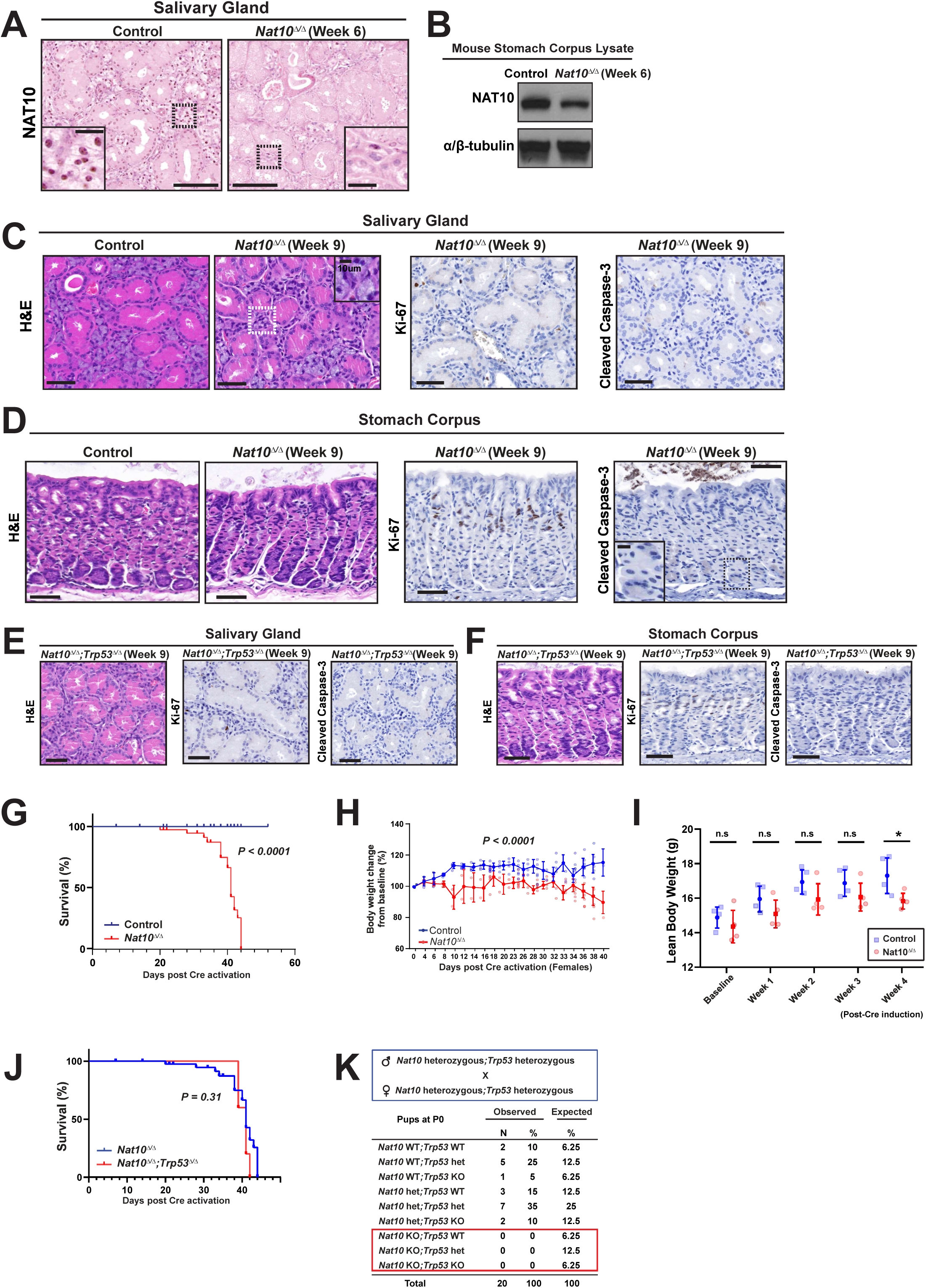
Loss of NAT10 yields a wide range of responses in various organs. (A&B) Cre induction in *Nat10^flox/flox^; Mist1^CreERT2/+^*mice results in a rapid reduction in NAT10 protein expression in acinar cells of the submandibular salivary gland (A; scale bars: 100 μm, inset: 20 μm) and stomach corpus (B), as shown by IHC and western blot, respectively. (C-F) Acinar cells of the salivary glands did not show consistent change in the morphology (C, left; scale bars: 50 μm, inset: 10 μm), increase in proliferation index (C, middle), or apoptosis (C, left). Likewise, chief cells at the base of the stomach did not show significant changes in cell volume or death phenotype until 12 weeks after Cre induction (D, scale bars: 50 μm, inset: 10 μm). Similarly, no significant differences in the morphologic, proliferative or apoptotic indices were observed in salivary gland acinar cells (E, scale bar: 50 μm) or stomach chief cells (F, scale bar: 50 μm) with p53 deletion (*Nat10*^Δ*/*Δ^*; Trp53*^Δ*/*Δ^). (G-K) Deletion of p53 does not rescue the lethality of mice deficient in NAT10. (G-J) *Nat10^flox/flox^; CAGG-Cre^CreERTM^* mice become moribund approximately 6 weeks after Cre recombination (G), with eventual loss of weight (H) and decrease in lean body weight (I). Concomitant loss of p53 in *Nat10*-deficient mice (*Nat10^flox/flox^; Trp53^flox/flox^ CAGG-Cre^CreERTM^*) does not rescue the mice from lethality (J). Similarly, embryonic lethality of *NAT10^−/−^* mice is not rescued by knocking out the *Trp53* gene, as evidenced by the lack of *Nat10^−/−^;Trp53^−/−^* (double-knockout) pups born from *Nat10^+/−^;Trp53^+/−^* Nat10^+/−^;Trp53^+/−^* breeding scheme (K).

We next asked whether global, irreversible deletion of NAT10 affects adult mice. For this purpose, we again used *Nat10^flox/flox^; CAGG-Cre^ERTM^* (*Nat10*^Δ*/*Δ^) mice, where we could induce universal Cre recombination in a temporally controlled manner. We noticed all of the *Nat10*^Δ*/*Δ^ mice became moribund, requiring euthanasia at a median of 41 days post-Cre induction (Fig. 7G), while all of the control mice survived (log-rank test, *P* < 0.0001). Although we are not able to precisely define the expression level of NAT10 in each organ and thus cannot determine the organ(s) influencing its demise, we noticed substantial weight loss before *Nat10*^Δ*/*Δ^ mice became moribund that was not noted in the controls (Fig. 7H, *P < 0.0001*), in addition to a significant decrease in lean body weight (Fig. 7I). As the consumption of chow did not differ between the control and *Nat10*^Δ*/*Δ^ mice for three weeks after Cre induction as determined by Comprehensive Lab Animal Monitoring System (CLAMS; week 1 to 2: 10.68 ± 1.28 vs. 10.17 ± 0.53, *P = 0.34*; week 3 to 4: 10.79 ± 1.85 vs. 9.71 ± 0.85, *P = 0.33*), inability to generate energy might have played a role.

We next sought to determine whether the *Nat10*^Δ*/*Δ^ death phenotype could be rescued by deleting p53. Such rescue has been noted in other ribosomopathic mouse models (Jones et al., 2008; Panić et al., 2006; Stadanlick et al., 2011), as well as in our pancreatic acinar cell studies. Interestingly, the survival of *Nat10^flox/flox^; Trp53^flox/flox^; CAGG-Cre^ERTM^ (Nat10*^Δ*/*Δ^*; Trp53*^Δ*/*Δ^) mice did not differ from that of the *Nat10*^Δ*/*Δ^ mice (Fig. 7J, median survival 41 days vs. 41 days; P = 0.31).

Therefore, the deletion of p53 did not rescue the mortality of *Nat10*^Δ*/*Δ^ mice, which indicates that RiBi itself plays a p53-independent role in mouse survival. Finally, we asked if p53 loss could rescue the phenotype of the *Nat10*^Δ*/*Δ^ mice during development. *Nat10* homozygous knockout mice were reported to be embryonically lethal before E14.5 (Balmus et al., 2018), and indeed we never found any *Nat10* homozygous knockout pups from the *Nat10* heterozygous X *Nat10* heterozygous breeding pairs. We therefore set up a *Nat10* heterozygous; *Trp53* heterozygous X *Nat10* heterozygous; *Trp53* heterozygous breeder cage and examined the pups to determine if knocking out p53 could rescue the pups from embryonic lethality. Out of 20 pups examined, we found no *Nat10*^-/-^ *Trp53*^-/-^ weanlings (Fig. 7K), consistent with inability of p53 loss to rescue NAT10 loss during development. In sum, stabilization of p53 is insufficient to rescue the animal from mortality, suggesting the importance of RiBi and translation itself in crucial tissues in adult and embryonic mice.

### NAT10 is required for pancreatic tumorigenesis

p53 has long been deemed a gatekeeper for cancer formation and progression. Given the action of p53-induced cell death and comparison with the dual action of NAT10 loss on cell function, we chose to interrogate whether NAT10, or more broadly, RiBi, is critical *in vivo* in long-lived cells that are recruited as cellular sources of solid tumors. We used a well-characterized pancreatic adenocarcinoma tumorigenesis model, the *Kras^G12D^* knock-in mouse model, which drives acinar cells into ADM, followed by pancreatic intraepithelial neoplasia (PanIN), then finally into invasive pancreatic ductal adenocarcinoma (PDAC). Within 3 months after Cre induction (Fig. 8A), we noticed multifocal PanIN lesions in the pancreas (Fig. 8B, left panels) with a high proliferation index (Fig. 8B, right) in the control mice (*Kras^G12D^ mice; Nat10^flox/+^; Kras^G12D/+^; ROSA26^LSLTdTomato/+^; Mist1^CreERT2/+^*). PanIN lesions were not only positive for ADM markers CK19 and SOX9 (Fig. 8C), as expected, but also were positive for TdTomato signal when lineage-traced, confirming their acinar cell origin. In contrast, we did not notice any PanIN lesions in any of the pancreases of the *Nat10*^Δ*/*Δ^; *Kras^G12D^*mice *(Nat10^flox/flox^; Kras^G12D/+^; ROSA26^LSLTdTomato/+^; Mist1^CreERT2/+^*, Fig. 8D), and the morphology of these quiescent acinar cells was identical to that of the *Nat10*^Δ*/*Δ^ mice. Likewise, nearly all CK19/SOX9-positive cells were negative for TdTomato, indicating they were normal ductal cells and not acinar cells having undergone ADM (Fig. 8E). PanIN lesions in the control mice included the expected Alcian-blue positive mucinous foci (Fig. 8F), the area of which was significantly higher in *Kras^G12D^* mice compared to *Nat10*^Δ*/*Δ^; *Kras^G12D^* mice (23.6 ± 12.7 % vs. 0 ± 0 % of total cross-sectional area, *P* = 0.0098, Fig. 8G). Thus, NAT10, and by extension RiBi, is required for the development of PanIN lesions.

**Fig. 8.**
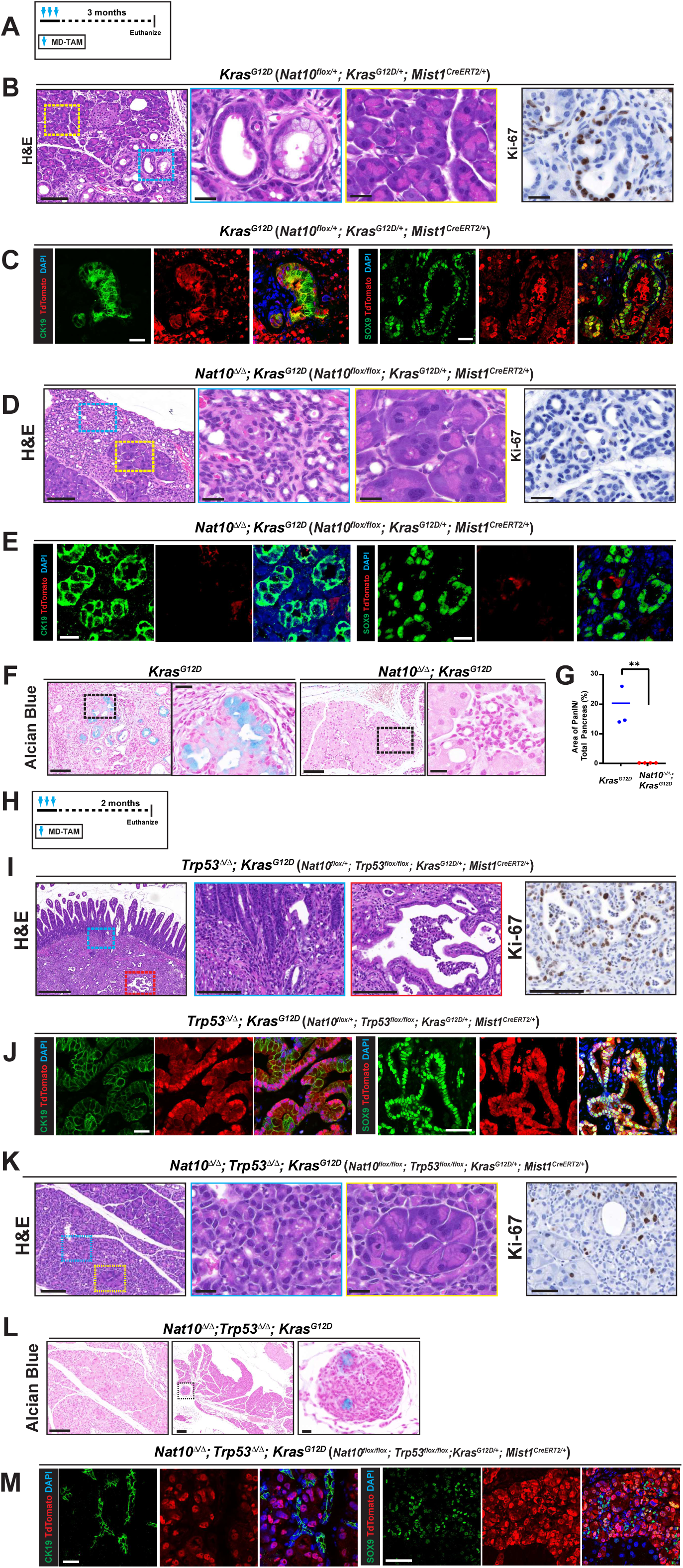
NAT10 is required for PanIN and PDAC formation. Mice with conditional expression of mutant Kras (*Kras^G12D^*) were generated following the *Mist1^CreERT2^* activation scheme using medium dose tamoxifen (MD-TAM, A). Control mice developed multifocal PanIN lesions (B, left panels, blue box, scale bars: 100 μm, insert 20 μm), which were positive for Ki-67 (B, right panel; scale bars: 100 μm, insert 20 μm). These PanIN lesions were also positive for CK19 (C, left panels), SOX9 (C, right panels), and TdTomato, demonstrating their acinar cell origin (C; scale bars: 20 μm). In contrast, shrunken cells with scant cytoplasm dominated the pancreas of *Nat10*^Δ*/*Δ^*; Kras^G12D^* mice without evidence of PanIN lesions on H&E staining (D, left panels; scale bar: 100 μm, insets 20 μm). Shrunken cells did not enter the cell cycle (D, right panel; scale bar: 20 μm). In these mice, CK19 or SOX9 positive cells lacked TdTomato signal, suggesting those are not PanINs but remaining ductal cells (E; scale bar: 10 μm). Likewise, the percentage of PanIN area in *Kras^G12D^* pancreas, which contained Alcian blue-positive mucinous glands (F, scale bar: 100 μm, inset 20 μ), was significantly higher than in *Nat10*^Δ*/*Δ^ pancreas (G). (H-M) NAT10 is required for PDAC formation in the absence of p53. Mice with p53 knocked out and expressing the Kras mutant (*Trp53*^Δ*/*Δ^*; Kras^G12D^*) were generated following the Cre activation scheme using tamoxifen (H). These mice showed PDAC invading adjacent organs such as the small intestine (I, left panels; scale bars: 500 μm, insets 100 μm) with high proliferation index (I, right panel; scale bar: 100 μm). PDAC lesions were positive for CK19 (J, left panels; scale bar: 20 μm), SOX9 (J, right panels; scale bar: 50 μm), and TdTomato, demonstrating their acinar cell origin. In contrast, mice deficient in NAT10 and p53 with the *Kras^G12D^* mutation (*Nat10*^Δ*/*Δ^*; Trp53*^Δ*/*Δ^*; Kras^G12D^*) did not develop invasive PDAC lesions, as shown by H&E staining (K, left panels; scale bars: 100 μm, inset 20 μm), although the cells readily re-entered the cell cycle (K, right panel; scale bar: 50 μm). Alcian-blue positive PanIN lesions were largely absent (L, left panel; scale bar: 100 μm), and only rarely noted (L, two right panels, scale bars: 200 μm, inset 20 μm) in *Nat10*^Δ*/*Δ^*; Trp53*^Δ*/*Δ^*; Kras^G12D^* mice. Likewise, the vast majority of CK19 or SOX9-positive cells in *Nat10*^Δ*/*Δ^*; Trp53*^Δ*/*Δ^*; Kras^G12D^* were of ductal origin, as shown by the negative TdTomato staining (M; scale bars: 20 μm (left), 50 μm (right).

We next asked if NAT10 was required for tumorigenesis in the p53 null context to better reflect the actual pancreatic tumorigenesis process, which frequently shows mutations resulting in the loss of p53 function (Morton et al., 2010). To this end, we tested *Trp53*^Δ*/*Δ^*; Kras^G12D^* mice (*Nat10^flox/+^; Kras^G12D/+^; Trp53^flox/flox^; Mist1^CreERT2/+^*), which, within two months after Cre induction (Fig. 8H), developed PDAC invading adjacent small intestine (Fig. 8I, left panels) and with a high proliferative index (Fig. 8I, right). PDACs were positive for CK19 and SOX9, as well as TdTomato lineage-tracing (Fig. 8J), consistent with acinar cell origin of the tumor cells. In contrast, the *Nat10*^Δ*/*Δ^*; Trp53*^Δ*/*Δ^*; Kras^G12D^* mice (*Nat10^flox/flox^; Kras^G12D+/^; Trp53^flox/flox^; Mist1^CreERT2/+^*) showed similar morphologic features (Fig. 8K, left) and proliferative index (Fig. 8K, right) comparable with *Nat10*^Δ*/*Δ^*; Trp53*^Δ*/*Δ^ acinar cells, without obvious evidence of tumor, and only a few scattered Alcian Blue positive PanIN lesions (Fig. 8M, n=8) present two months after Cre induction. Recombined acinar cells were still alive and positive for TdTomato (Fig.8L). However, they were mostly negative for CK19 (Fig. 8M, left) and SOX9 (Fig. 8M, right), indicating that RiBi and translation are still required for sequential ADM-PanIN-PDAC formation, even in the absence of p53.

Because the first step in tumorigenesis is the paligenotic conversion of acinar cells to ADM, marked by induction of SOX9 and CK19, the results are consistent with absolute requirement for NAT10 and RiBi, the data indicate loss of RiBi blocks tumorigenesis because it blocks the the initial paligenosis required to trigger tumorigenesis.

## DISCUSSION

In this study, we employed a novel NAT10 conditional knock-out mouse model, as well as various Cre lines, lineage tracing, and *ex vivo* culture models to determine how mature differentiated cells respond to defects in RiBi. Our data revealed a unique susceptibility of pancreatic acinar cells to loss of NAT10, resulting in p53-induced apoptotic cell death as well as structural and functional cell rearrangement that was not influenced by p53. Both phenotypes occurred only after a prolonged latency. The findings herein highlight the added value of using a cell type-specific approach that targets specific biological processes (e.g., RiBi or translation) and structures (e.g., ribosomes) in the study of cell behavior *in vivo* in multicellular organisms.

To date, the influence of RiBi inhibition on cell fate has been investigated overwhelmingly *in vitro.* Loss of RiBi has been shown consistently to cause rapid induction of cell cycle arrest, with resulting senescence or death. Loss of *Bop1*, a nucleolar protein involved in rRNA processing and ribosome assembly, resulted in cell cycle arrest through p53 stabilization in a murine-derived cell line (Pestov et al., 2001). Treatment with BMH-21, a pol-I inhibitor that stalls RiBi, induced p53-dependent inhibition of cell growth and blocked tumor growth in cancer-derived cell lines. The RiBi blockade and p53 induction occurred without DNA damage (Peltonen et al., 2010). CX-5461, a TOP2 inhibitor with indirect pol-I inhibition function, induced p53-stabilization and subsequent apoptosis in cancer cell lines (Bywater et al., 2012). Such *in vitro* systems are beneficial for understanding the importance of RiBi in general cellular behavior; however, tissue culture systems are also limited in that they generally do not model the behavior of non-dividing cells. Furthermore, in non-dividing cells, particularly in long-lived, terminally differentiated cells that must perform translation-rich activities like regulated secretion of proteins, the census of ribosomes would be expected to accumulate and be maintained at much higher levels than in cells that divide frequently, while each cell division would reduce the pool of ribosomes by about one half in resulting progeny cells. Given that the vast majority of cells in most multicellular organisms are differentiated and non-dividing (Cho et al., 2024), *in vitro* systems may be modeling the importance of RiBi in only a small fraction of actual cells in living tissue. In the current study, we focused on effects of blocking RiBi *in vivo*, targeting the long-lived, acinar cells in the exocrine pancreas of mice.

Pancreatic acinar cells are archetypical of differentiated cell types found in multicellular organisms (Cho et al., 2024). They are characterized by long half-life, large size with disproportionately enriched rough ER and ribosomal contents, and lack of proliferation (quiescence) at homeostasis. To date, our understanding of the response to aberrant RiBi and translation in the pancreas has been mostly limited to studies on developmental aspects such as Shwachman-Diamond syndrome (SDS) or Shwachman-Diamond-like syndromes, which encompass congenital ribosomopathies that share similar phenotypes. In these cases, pancreatic insufficiency along with various degrees of involvement of other organs, occurs from mutations in genes related to RiBi, translation, or signal recognition particle-mediated ribosome translocation to rough ER, including *SBDS* (Boocock et al., 2003; Tourlakis et al., 2012; Venkatasubramani & Mayer, 2008), *SRP54* (Bellanné-Chantelot et al., 2018; Carapito et al., 2017), *SRPRA* (Linder et al., 2023), and *EFL1* (Tan et al., 2019). Although these congenital ribosomopathies provide irrefutable evidence about the susceptibility of pancreatic acinar cells to the loss of RiBi and translation, patient data and mouse models for these germline-mutation-induced disease states provide only limited insight into how this fully differentiated cell type responds to defects in RiBi and translation in an adult, non-developmental, context. Monitoring sequential changes that occur in response to RiBi blockade in specific cell types requires models that permit: a) cell-type specificity; b) the ability to manipulate a RiBi-regulating gene or genes of interest in a temporally controllable environment; c) tools to read out results *in vivo*. The results of our current study, in which we present a model system to study all these aspects of RiBi in a single, differentiated cell type *in vivo*, have very minimal precedent for discussion within the context of existing literature.

It is currently unclear whether inhibition of ribosomes and/or translation universally results in p53-stabilization and subsequent cell death, or merely inhibits proliferation with defective translation. In particular, the question of how p53 correlates with ribosomal function has not been studied in depth in cells *in vivo*. Our results imply that the cellular response to the loss of component(s) required for RiBi and translation in tissue should be interpreted within a cell type- and tissue-specific context, as noted by the different responses observed in the acinar cells of the pancreas and salivary gland, as well as in the chief cells of the stomach. Furthermore, our results show that p53 is required to halt cell division and induce cell death when RiBi and translation are compromised because loss of p53 rescued the cell death phenotype in acinar cells lacking *Nat10*. In fact, cells lacking *Nat10* and *Trp53* not only survived but could proliferate (presumably *Nat10*^Δ*/*Δ^;*Trp53*^Δ*/*Δ^ cells are induced to proliferate to try to restore perceived deficiency in enzyme-secreting cells).

However, p53 is not involved in the loss of physiological function in acinar cells caused by compromised RiBi, as *Nat10*^Δ*/*Δ^;*Trp53*^Δ*/*Δ^ cells shrank and lost digestive-enzyme cargo similar to cells lacking only *Nat10*. We also observed that loss of p53 alone in an uninjured pancreas has no noticeable effect in this quiescent organ, as others have also noted (Jacks et al., 1994). Rather, p53 is required only when these cells, which are otherwise quiescent, re-enter the cell cycle (Marcel et al., 2013; Miao et al., 2020; Rosenfeldt et al., 2013; Yang et al., 2014). It is surprising not so much that loss of NAT10 is registered by acinar cells eventually as an “injury” that induces p53 but that recognizing defective RiBi as an injury takes acinar cells several weeks. Thus, a ‘tipping point’ or threshold of some sort appears to exist before the cells react to loss of RiBi. What dictates this tipping point is still unclear. Perhaps a critical number of ribosomes exist in a certain type of cell to execute that cell’s physiological function, such that below that threshold, injury is sensed. Alternatively, and not mutually exclusively, it is the lack of overall tissue function (i.e., loss of digestive enzyme secretion) that releases cell-exogenous signals to stimulate proliferation. The questions of which cell-autonomous or exogenous signals read eventual loss of RiBi and how those signals are related to p53 in certain types of cells (but not others) should be further investigated. Finally, the loss of 18S rRNA in *Nat10*^Δ/Δ^ acinar cells was not reversed by the concomitant loss of p53 (Fig. 6J), indicating that p53 does not affect rRNA biogenesis *in vivo*. All the data herein indicate that p53 does respond to RiBi, but the interactions between RiBi and p53 are complex and contextual *in vivo* in non-malignant cells.

We also investigated the role for specific ribosomal proteins in response to decreased RiBi and in correlation with p53. Both RPL5 and RPL11, which influence p53 stabilization through inhibition of its interaction with MDM2 (Lohrum et al., 2003), showed a dramatic change in their localization in pancreatic acinar cells, while their absolute levels decreased. This is similar to previous reports that the disruption of the nucleolus results in the RPL5/RPL11/5S rRNA complex, a.k.a., 5S ribonucleoprotein particle (RNP) interacting with MDM2, resulting in p53 stabilization under nucleolar stress conditions (Bursać et al., 2012; Pestov et al., 2001; Rubbi & Milner, 2003). However, it should be noted that deletion of NAT10 does not disrupt the nucleolus *in vivo* in acinar cells and directly affects the maturation of only 18S rRNA (and not 28S, 5.8S, and 5S rRNA). Therefore, the ribosomal localization changes we observe differ from ‘classical’ nucleolar stress induced by pol-I inhibition (though, again, the caveat being nucleolar stress has been defined mostly in tissue culture). Although NAT10 is required for 18S rRNA synthesis, we noted that not only ribosomal small subunit (SSU) but also ribosomal small subunit (LSU) components were degraded in *Nat10*^Δ*/*Δ^ cells. At the same time, nuclear translocation of 5S RNP components, which are also part of the LSU, occurred when *Nat10* was deleted. This suggests that pancreatic acinar cells might sense the discrepancy between LSU and SSU in the cytoplasm or the imbalance of 5S RNP relative to other mature ribosomal components. Somehow this imbalance may trigger 5S RNP to cause p53 stabilization via MDM2 sequestration. It is possible that part of the accumulation of RPL5 and RPL11 in the nucleoplasm may be due to a failure in their export to the cytoplasm; however, as they are translated in the cytoplasm, there must be at least some rerouting into the nucleus. It is not clear what fraction of the translocating LSU components originate from disassembly of existing ribosomes in the cytoplasm vs from a pool of translated proteins that have never been incorporated into ribosomes.

The p53-dependent cell-cycle blockage we report here is in line with previously reported impairment of mitogenic signal-associated post-hepatectomy regeneration in a mouse model with conditionally deleted *Rps6* (an integral component of SSU). Interestingly, in that report, unaltered translation in the refed liver after fasting was observed (Volarevic et al., 2000), which contrasts with the phenotype we observed in pancreatic acinar cells. It is important to note that the latency period was not precisely documented in the previous report, and testing the phenotype after prolonged latency may help rule out the possibility that low levels of mature ribosomes are present and sufficient to mediate normal translation functions, as we hypothesize helps account for the latency in our current study. This question is particularly relevant given the fact that hepatocytes are, like acinar cells, differentiated, long-lived, quiescent cells. Further studies should determine whether RiBi is active in this organ and the rate of extant cytosolic ribosome turnover in the absence of RiBi.

Pancreatic acinar cells are known to undergo acinar-to-ductal metaplasia (ADM), where a loss of their organelles leads to a morphological change that resembles pancreatic ductal cells (Giroux & Rustgi, 2017; Ma et al., 2022; Mills & Sansom, 2015). Acinar cells undergo ADM by paligenosis, an evolutionarily conserved cellular program that consists of three stages occurring in differentiated cells recruited to divide by tissue injury: autodegradation → metaplastic gene expression → p53 licensing of cell cycle reentry (Brown et al., 2022; Miao et al., 2020; Radyk et al., 2021; Willet et al., 2018). We noted that *Nat10*^Δ*/*Δ^ acinar cells did not undergo paligenosis; transmission EM did not reveal obvious induction of autophagosomes or lysosomes, and *Nat10*^Δ*/*Δ^ cells were shrunken but still negative for SOX9, which is a marker of the second phase of paligenosis. However, paligenosis is an acute process that occurs over the course of days in response to external stimuli, and we observe dynamic regulation of ribosomes during the process (unpublished data). We anticipate that *Nat10*^Δ*/*Δ^ acinar cells may help us understand how pancreatic acinar, or other types of differentiated, cells manage RiBi and ribosomes, in injury. Likely RiBi will be a key component of paligenosis that *Nat10*^Δ*/*Δ^ cells may help us understand. It will be especially interesting to further examine whether nucleoplasmic localization of 5S RNP components occur during the normal p53 licensing (stage 2 to 3) step during paligenosis.

PDAC is the seventh most frequent source of cancer-related death worldwide, with a poor prognosis in advanced stages (Bengtsson et al., 2020; Ferreira et al., 2017; Rawla et al., 2019; Sung et al., 2021). It typically arises in cells that have mutations in the an oncogenic driver gene, *Kras*, along with well-known additional mutations (e.g., in genes encoding p53, p16, SMAD4, PTEN), which all conspire to initiate and propagate the precancerous lesion, PanIN (Grimont et al., 2022). Secretory acinar cells of the pancreas appear to be one of the principal cells of origin for PanIN and PDAC (Kopp et al., 2012), facilitated by repeated bouts of acute pancreatitis or chronic pancreatitis that induce ADM. Therefore, understanding how acinar cells work at homeostasis is crucial for understanding origins of PDAC and, potentially for evaluating the efficacy and side-effects of PDAC treatments that might target ribosomes. As cancer is a disease unique to multicellular organisms (Cho et al., 2024), therapeutic development requires a comprehensive understanding of the characteristics of the cell types from which cancer originates. Here, we show that targeting a component of RiBi might work as a tumor suppression not just by targeting p53, as many other cancers do (via triggering DNA damage response) but also by directly inhibiting protein translation in a p53-independent fashion. This provides support that targeting RiBi in conjunction with other therapeutic agents, especially in the context of p53 mutations, might be parallel and hence complimentary approaches. Furthermore, the surprising latency in response to RiBi and translation block, as well as the finding that not all cells *in vivo* in tissue respond in the same manner, both support the idea that RiBi and translation can potentially be targeted chemotherapeutically for organ-specific or cell type-specific treatments. Side effects on normal tissue may be limited if treatment duration is limited.

In sum, this is the first comprehensive and rigorous evaluation of NAT10 function in RiBi and translation in long-lived adult differentiated cell types *in vivo* and *ex vivo*. The study revealed varying responses among different cell types in an adult organism. The methods we present here can be adapted to assess effects of loss of RiBi and translation in other types of differentiated cells. Such better understanding will help us decipher roles of p53-RiBi interactions *in vivo* in homeostasis, which may also help us develop novel chemotherapeutic approaches with potentially more targeted effects.

## METHODS

### Mice

All animal experiments were performed according to protocols approved by the Washington University School of Medicine and Baylor College of Medicine Animal Studies Committees. The *Nat10* floxed mouse was created using reagents designed, validated, and genotyped by the Genome Engineering & Stem Center (GESC@MGI) at Washington University in St. Louis. Briefly, gRNAs were designed to mediate insertion of loxP sites into introns 1 and 3, effectively floxing the first two coding exons. A single-stranded oligodeoxynucleotide (ssODN) with loxP flanked by short homology arms in the CRISPR/Cas9 cleavage site was used as the donor template with each gRNA (see Supplementary Table 1 for the list of gRNAs and ssODNs). The gRNA/Cas9 protein complexes, together with ssODNs which were ordered as ultramers from ITD, were transfected into N2A cells and validated by NGS for efficient insertion of loxP at Cas9 cleavage sites before use in mice. Live births were genotyped using NGS. Mouse work was performed at the Transgenic, Knockout, and Microinjection Core at Washington University in St. Louis. gRNA/Cas9 complexes, along with ssODNs, were electroporated into single cell embryos and transferred into pseudopregnant recipients to generate founder animals. Six founder mice were born, mated to generate F1 progeny, and further bred to various Cre alleles to examine the phenotype of *Nat10* loss *in vivo*. 6- to 20-week-old mice were used in experiments, with salivary gland acinar cell phenotype assessed only in male mice for consistency. Medium-dose tamoxifen (3 mg/20 g mouse body weight, intraperitoneal injections) was administered for three (for pancreatic tumorigenesis study) or five consecutive days to induce recombination of Cre in the pancreas or salivary gland in *Mist1^CreERT2^* mice or activate Cre in *CAGG-Cre^ERTM^* mice. Low-dose tamoxifen (1 mg/20 g mouse body weight, intraperitoneal injections) was administered for 7 consecutive days to induce recombination of Cre in the stomach. Tamoxifen Toronto Research Chemicals, Inc, Toronto, Canada) was prepared in a 10% ethanol and 90% sunflower oil solution by sonication as previously described (Saenz et al., 2016). For the *in vivo* BMH-21 treatment, BMH-21 was dissolved in citrate phosphate buffer (1 mg/200 uL/20 g) and was given intraperitoneally once a day for 5 days or twice a day for 3 days. Control mice received citrate phosphate buffer alone. For measurement of whole-body fat mass, lean tissue mass, free fluid, and total body water, the nuclear magnetic resonance system EchoMRI-900™ (Echo Medical System, Houston, TX) was used. When examining food intake, mice were maintained on a high-fat diet and housed at room temperature in Comprehensive Lab Animal Monitoring System Home Cages (CLAMS-HC, Columbus Instruments). O_2_ consumption, CO_2_ emission, energy expenditure, food and water intake, and activity were measured for seven days at BCM Mouse Metabolic Phenotyping Core.

### Cell culture, transfection, drug treatment, and cell viability assay

AGS and LS-174T cells were cultured in RPMI-1640 medium with 10% fetal bovine serum and Primocin (100 µg/mL). All cells were maintained at 37 °C in a 5% CO_2_ humidified atmosphere. Transient knockdown of NAT10 was carried out using two siRNAs against NAT10 (SASI_Hs01_00215377; siNAT10#1, and SASI_Hs02_00357064; siNAT10#2, Sigma). MISSION® siRNA Universal Negative Control (Sigma) was used as a negative control. Briefly, AGS cells were seeded in 6-well flat bottom plates at a density of 1 x 10^5^ cells per well, transfected with siRNA in Opti-MEM™ (Gibco) using Lipofectamine 2000 (Invitrogen) according to the manufacturer’s protocol, and incubated for 24 to 72 hours. Cell viability was assessed using alamarBlue™ assays (Invitrogen). 1 to 10 µM of BMH-21, or CX-5461 (100 nM), or diluents were added in triplicate, and the cells were incubated for 24 hours. The alamarBlue reagent was added to each sample at a 10% final concentration and incubated for 3 hours. Viability was analyzed by measuring absorbance at 570 nm and 600 nm, and the percentage of alamarBlue reduction was calculated. The values of reduction were corrected for the background values of media without cells.

### Western blot, immunoprecipitation, and subcellular fractionation

To extract protein, cells were lysed in RIPA buffer with Halt™ Protease and Phosphatase Inhibitor Cocktail (Thermo). When using mouse tissues, approximately 35 to 70 μg of tissue lysate were lysed in T-PER Tissue Protein Extraction Reagent buffer (Thermo). Lysates were clarified by centrifugation at 16,000 × g for 10 minutes at 4 °C to remove cell debris. Proteins were separated using a 4-12% gradient SDS-PAGE gel and transferred to a 0.2 or 0.45 µm nitrocellulose or PVDF membrane, blocked with 3% Bovine serum albumin (BSA) in tris-buffered saline with 0.1% Tween-20 (TBS-T), and then incubated for 16 hours at 4 °C with primary antibodies (see Supplementary Table 2 for the list of antibodies). The membranes were washed three times in TBS-T, incubated for 1 hour at room temperature with HRP-conjugated secondary antibodies in 5% nonfat dairy milk or 3% BSA in TBS-T. Secondary antibody signals were imaged and detected using the SuperSignal West Pico PLUS Chemiluminescent Substrate Kit (Thermo). Protein signal intensities were normalized against a β-actin or alpha/beta tubulin loading control for each sample.

For immunoprecipitation, lysates from cells or tissues were prepared using Pierce IP lysis buffer (Thermo) with protease/phosphatase inhibitors. 50 μL of Protein A beads were washed with wash buffer (phosphate-buffered saline with 500 mM NaCl and 0.1% Tween 20) and incubated with either anti-NAT10 antibody or normal rabbit IgG (negative control; Cell Signaling) for 20 minutes at room temperature with rotation, washed three times in wash buffer, and incubated overnight at 4 °C with rotation with the tissue or cell lysate. Next, the beads were washed five times with wash buffer and were subsequently used for mass spectrometry or western blotting experiments. The quantitative value (normalized total spectra) of proteins enriched in NAT10 or IgG pulldown lysates from mass spectrometry data were exported from the Scaffold viewer (version 5) using the criteria of protein threshold of 95%, minimum number of peptides of 2, and peptide threshold of 1% FDR. Proteins having more than 2-fold enrichment in NAT10 pulldown compared to IgG were further analyzed for gene ontology using DAVID (Dennis et al., 2003). GO terms under the biological process category were plotted using Rstudio/R with the GO terms, fold enrichment, and FDR values from the DAVID results. Subcellular fractionation was performed following the published protocol. (Lam & Lamond, 2006)

### Pancreas 3D organoid generation and 2D culture

Mouse pancreas organoids were generated with minor modifications to a widely used protocol (Broutier et al., 2016). Following euthanasia, the mouse pancreas was dissected and finely chopped into 0.5 mm^3^ sections after being rinsed three times in wash medium (DMEM high glucose + 1% FBS + sodium pyruvate). The supernatant was discarded, and the pancreatic fragments were transferred to a 50 mL tube.

Following double rinsing with wash medium, the samples were digested at 37 °C with a solution containing DNase I, collagenase, and dispase while being agitated at 200-250 RPM. After digestion, the supernatant was transferred to a 15 mL tube and placed on ice. The sample was subsequently centrifuged twice at 800 RPM in ice-cold wash medium at 8 °C for 5 minutes after passing through a 70 μm cell strainer. The cells were washed twice with 10 mL of cold basal medium (Advanced DMEM/F12+ GlutaMAX+ HEPES), resuspended in 50 μL of Matrigel, and then transferred to a pre-warmed 24-well plate. 500 uL of pre-warmed conditioned medium (Advanced DMEM/F12+ R-Spondin/Noggin conditioned medium) supplemented with HEPES, GlutaMAX, N-acetylcystein, Nicotinamide, EGF, FGF10, and Gastrin were added to each well. The medium was changed every 3 days, and the images were taken with a Lionheart FX Automated Microscope and Live Cell Imager (BioTek). Pancreatic acinar cells were also grown as two-dimensional (2D) culture on collagen-coated plates following the previously published protocol (Gout et al., 2013). Briefly, pancreas was prepared in a similar manner as above and digested with a collagenase 1A solution (0.25 mg/mL) with HEPES (10 mM) and Soybean Trypsin inhibitor (SBTI, 1 mg/mL) at 37 °C. The samples were subsequently centrifuged at 450 × g in ice-cold wash medium at 4 °C for 3 minutes, repeated five times, and passed through a 100 μm cell strainer in Waymouth medium supplemented with Primocin, 2.5% FBS, EGF (25 ng/mL), and 0.25 mg/mL of SBTI. Isolated acini were initially seeded in non-coated 6-well plates and subsequently transferred to a collagen coated plate.

### Imaging and tissue analysis

Mouse tissue was fixed in 10% neutral buffered formalin for 16 hours at 4 °C, washed in PBS, and transferred to 70% ethanol. Stomach samples were immobilized in 3% agar. All samples were embedded in paraffin and sectioned at 5 to 7 µm thickness. For IHC, sections were deparaffinized in Histo-Clear (National Diagnostics), rehydrated through a series of ethanol washes, and quenched for endogenous peroxidase for 15 minutes. Heat-induced epitope retrieval was performed in Sodium Citrate buffer (pH 6.0). Samples were subsequently blocked with 5% normal serum in 0.15% Triton X-100 for 1 hour in room temperature and incubated with primary antibodies for 16 hours at 4 °C. After three PBS washes, samples were incubated for 1 hour with biotin-conjugated secondary antibodies, followed by incubation with avidin/biotin complex (ABC, Vector Laboratories) for 1 hour at room temperature. Sections were developed with the DAB Substrate Kit and counterstained with Hematoxylin and/or Eosin prior to mounting. For IF staining, sections were deparaffinized in xylene, rehydrated in isopropanol, and underwent heat-induced epitope retrieval in sodium citrate buffer (pH 6.0). Samples were then blocked with 1% BSA in 0.15% Triton X-100 for 1 hour at room temperature and incubated overnight with primary antibodies. Primary antibodies were detected with Alexa Fluor-conjugated secondary antibodies (Invitrogen) and mounted in ProLong Gold Antifade Mountant with 4′6-diamidino-2-phenylindole (DAPI) visualize nuclei. IF images were captured using AX R Confocal System with Eclipse Ti2-E Inverted Microscope (Nikon) or LSM880 Laser Scanning Confocal Microscope (Zeiss), ensuring consistent exposure times across samples for comparison. Alcian blue staining was carried out using Alcian Blue (pH 2.5) Stain Kit (Vector Laboratory) following manufacturer’s instructions.

IF of cells was performed on cultures grown on coverslips in 12-well culture plates. Cells were washed with PBS, fixed with 4% paraformaldehyde, washed three times in PBS, permeabilized with 0.2% Triton X-100 for 15 minutes at room temperature, and blocked with 1% BSA in 0.15% triton X-100 for 1 hour at room temperature.

Cells were then incubated overnight with primary antibodies. After three PBS washes, samples were incubated for 1 hour with Alexa Fluor-conjugated secondary antibodies (Invitrogen) and mounted onto slides with ProLong Gold Antifade Mountant with DAPI.

### Northern blot, ac4C Immuno-Northern blot, RNA gel electrophoresis, ac4C sequencing

For Northern blot and ac4C Immuno-Northern blot, 5 µg of RNA was extracted from whole kidney using TRIzol™ Reagent. RNA was loaded and separated on an agarose-formaldehyde gel following the published protocol (Mansour & Pestov, 2013). This method of RNA gel electrophoresis was used throughout the manuscript to determine the 18S/28S rRNA ratio. Gel images were captured using a ChemiDoc™ Imager (Bio-Rad) and transferred to Hybond-N+ positively charged Nylon membranes (Amersham Life Sciences) using a downward capillary transfer method. Northern blots were performed using the Northern blot starter kit (Roche), following the manufacturer’s protocol. A digitonin-labeled RNA probe with a complementary sequence to ITS-1 was generated from IDT (sequence: 5’-ttctctcacctcactccagacacctcgctccaca). Prehybridization was performed for 30 minutes, followed by hybridization at 68 °C for 16 hours with gentle agitation. The blots were incubated with an anti-Digoxigenin antibody and bands were visualized using a CDP-Star chemiluminescence reagent. ac4C Immuno-Northern blot and ac4C sequencing were carried out following the previously published methods (Bryson et al., 2020; Thalalla Gamage et al., 2021).

### 5-Ethynyl uridine (EU) pulse-chase experiment

Mice were pulsed with 2 mg of 5-Ethynyl uridine (EU) dissolved in 200 µL of PBS via intraperitoneal injection. After the given time, the mice were euthanized, and the pancreas tissue was snap frozen in O.C.T. compound using 2-methylbutane and liquid nitrogen, and sectioned using a cryotome at 9 µm thickness. The slides with tissue sections were fixed with 4% PFA for 10 minutes. To visulalize nascent RNA through the “click” reaction, the Click-iT™ RNA Alexa Fluor™ 594 Imaging Kit was used, following the manufacturer’s instructions.

### Sucrose gradient fractionation

Sucrose gradient fractionation was carried out with minor modifications to a published protocol (Blanc et al., 2023; Panda et al., 2017). Briefly, mouse kidney tissues were lysed in Pierce™ IP Lysis Buffer (Thermo) with Halt™ Protease and Phosphatase Inhibitor Cocktail (Thermo), 100 µg/mL cycloheximide, and 200 U of RNase OUT. Ribonucleoprotein complex was extracted using polysome extraction buffer (20 mM Tris-HCl pH 7.5, 100 mM KCl, 5 mM MgCl_2_, 0.5% NP-40, supplemented with 200 U of RNase OUT, Protease/phosphatase inhibitor, and 100 µg/mL cycloheximide). After centrifugation at 16,000 × g for 10 minutes at 4 °C, 1 mL of the lysates was loaded onto a 10%-50% sucrose gradient in a 14 mL polypropylene tube (Beckman Coulter). Ultracentrifugation was performed using a Beckman Optima LE-80K Ultracentrifuge Floor Centrifuge at 261,000 × g for 135 minutes at 4 °C. The centrifuge tubes were then removed from the bucket, placed on a tube holder, and pierced at the bottom with a 27 G needle. Multiple fractions (250 uL per fraction) were serially collected and labeled in approximately 40 x 0.5 mL microcentrifuge tubes and absorbance at 260 nm was measured for 2 μL aliquots of each fraction to determine the peaks (Blanc et al., 2023).

### RNA extraction and qRT-PCR

RNA was isolated using RNeasy mini kit (Qiagen) or Direct-zol RNA Miniprep Kits, and also underwent on-column DNase digestion per the manufacturer’s protocol. The quality of the mRNA was verified with a Nanodrop spectrophotometer (Thermo). 500 ng of RNA was reverse transcribed with the PrimeScript^TM^ RT Reagent kit (Takara) following the manufacturer protocol. Power SYBR Green master mix (Thermo) fluorescence was used to quantify the relative amplicon amounts of each gene (see Supplementary Table 3 for the list of the primers). For the Northern blot and RNA gel electrophoresis, we extracted RNA from the pancreas and kidney using TRIzol^TM^ reagent using the published protocol (Azevedo-Pouly et al., 2014).

### Transmission electron microscopy

Pancreas preparation and imaging for transmission electron microscopy (EM) were performed as previously described (Ramsey et al., 2007). Briefly, whole pancreas was collected, fixed overnight at 4 °C in modified Karnovsky’s fixative and sectioned. Pancreas tissue was processed and imaged at the Molecular Microbiology Imaging Facility, Department of Molecular Microbiology, Washington University in St. Louis. Nuclear circularity was measured by ImageJ (Schneider et al., 2012).

### Statistical analyses

Statistical analyses were performed using Prism 10 (GraphPad). In general, if datapoints were normally distributed, a t-test (for 2-sample assessments of error) was used to quantify the likelihood of true differences in means. Where distribution was non-normal, nonparametric tests were used. For comparisons between multiple groups, a one-way ANOVA with Dunnett’s multiple comparison post hoc test or Kruskal-Wallis test for non-normally distributed samples, was used to determine significance. A mixed-effects model followed by Sidak’s post-test for multiple comparison was used to compare the difference in weights of control and *Nat10^flox/flox^; CAGG-Cre^ERTM^* mice. Data were generally expressed as mean ± standard deviation (SD) unless, where statistical significance among multiple means was computed, in which case Standard Error of the Mean (SEM) was used. *P* < 0.05 was considered statistically significant for interpretation in the text.

## Acknowledgments

We thank Lillian Spatz, Ph.D. (Washington University in St. Louis), for assistance with the immunohistochemical staining. The expert technical assistance of Petra Erdmann-Gilmore, Dr. Yiling Mi, Alan Davis and Rose Connors is gratefully acknowledged. The proteomic experiments were performed at the Washington University Proteomics Shared Resource (WU-PSR), R. Reid Townsend MD.PhD., Director, Drs. Robert W. Sprung and Qiang Zhang, PhD., Co-Directors. The WU-PSR is supported in part by the WU Institute of Clinical and Translational Sciences (NCATS UL1 TR000448), the Mass Spectrometry Research Resource (NIGMS P41 GM103422) and the Siteman Comprehensive Cancer Center Support Grant (NCI P30 CA091842). Some of the microscopy and all tissue sectioning were performed by the AITAC of the Washington University Digestive Disease Research Core Center (DDRCC: P30DK052574) or by the Tissue Analysis and Molecular Imaging (TAMI) Core of the Texas Medical Center Digestive Disease Center (P30DK056338). We thank Wandy Beatty, Ph.D., with transmission electron microscopy studies at the Molecular Microbiology Imaging Facility at Washington University in St. Louis. We thank the GESC@MGI and Transgenic, Knockout and Microinjection Core at Washington University in St. Louis for creating the *Nat10* floxed mouse line. BCM Mouse Metabolism and Phenotyping Core is supported by NIH funds R01DK114356 and UM1HG006348.We also thank Robert Lawrence, Ph.D. for editorial support in review of the manuscript.

## Author Contributions

Study Concept and Design: CJC, TN, and JCM; Data Acquisition and Analysis: CJC, TN, AKR, YZH, ST, XL, STG, JLM and JCM; Drafting the manuscript: CJC, TN, ST, XL and JCM; Revisions to the manuscript: CJC, TN and JCM.

