## Supplementary table for "Inhibition of Ribosome Biogenesis *in vivo* Causes p53-Dependent Death and p53-Independent Dysfunction"

**Supplementary Table 1. List of guide RNA and single-stranded oligodeoxynucleotide sequences for generation of *Nat10* floxed mouse**

| Oligo | Sequence |
|---|---|
| gRNA for Intron 1 | ACTTTTAGGGATCCCCGTCC |
| gRNA for Intron 3 | ACTGTCTAGTACTGGTGAACNGG |
| ssODN for Intron 1 | a*t*ggcaaaaggaaagacttgccaggcagtggccgtatacacaagcagacccttcccaggaGGATCCATAACTTCGTATAGCATACATTATACGAAGTTATcggggatccctaaaagttcttctctggggctgaagagatggctcagtggttaagagca*c*t |
| ssODN for Intron 3 | g*a*tccctccaggaacagcgtttctccctttctcctctttagagagcttcttacaccagttGGATCCATAACTTCGTATAATGTATGCTATACGAAGTTATcaccagtactagacagttggtgattgtttacctctctc*t*c |

gRNA: Guide RNA

ssODN: Single-stranded oligodeoxynucleotide

*denotes a phosphorothioate bond

**Supplementary Table 2. List of primary nntibodies used in this study**

| Name | Company | Species | Cat. No. | Purpose | Dilution |
|---|---|---|---|---|---|
| NAT10 | Abcam | Rabbit | ab194297 | IB<br>IP<br>IHC | 1:1,000<br>5uL<br>1:250 |
| UBTF | Sigma | Rabbit | HPA006385 | IF | 1:200 |
| TCOF1 | Sigma | Rabbit | HPA038237 | IF | 1:200 |
| RPS6 | Cell Signaling | Rabbit | #2217 | IB | 1:2,000 |
| β-Actin | Santa Cruz | Mouse | sc-47778 | IB | 1:5,000 |
| ADAR1 | Santa Cruz | Mouse | sc-73408 | IB | 1:1,000 |
| HP1α | Cell Signaling | Rabbit | #2616 | IB | 1:2,000 |
| RPL11 | Abcam | Rabbit | ab79352 | IB<br>IF | 1:2,000<br>1:200 |
| RPL5 | Abcam | Rabbit | ab86863 | IB<br>IF | 1:2000<br>1:200 |
| RPL26 | Abcam | Goat | ab157111 | IF | 1:200 |
| HSPA5 | Cell Signaling | Rabbit | #3177 | IB | 1:1000 |
| SRP68 | Abcam | Rabbit | ab157120 | IB | 1:1000 |
| NPM1 | Abcam | Mouse | ab10530 | IF | 1:200 |
| α/β-tubulin | Cell Signaling | Rabbit | #2148 | IB | 1:2,000 |
| N4-acetylcytidine (ac4C) | Abcam | Rabbit | ab252215 | IB | 1:1,000 |
| CPA1 | R&D | Goat | AF2765 | IF | 1:200 |
| Amylase | Cell Signaling | Rabbit | # 3796 | IHC<br>IF | 1:500<br>1:500 |
| TdTomato | SicGen | Goat | AB8181 | IF | 1:200 |
| Cleaved caspase-3 (Asp175) | Cell Signaling | Rabbit | #9664 | IHC | 1:1,000 |
| SOX9 | Millipore | Rabbit | AB5535 | IHC<br>IF | 1:1,000<br>1:200 |
| p53 | Leica | Rabbit | CM5 | IHC<br>IB | 1:200<br>1:1000 |
| MDM2 | Abcam | Rabbit | ab259265 | IB<br>IF | 1:1000<br>1:200 |
| Ki-67 | Abcam | Rabbit | ab15580 | IHC | 1:200 |
| Ki-67 | Invitrogen | Rat | 14-5698-82 | IF | 1:500 |

| | | | | | |
|---|---|---|---|---|---|
| p-Histone H3 | Cell Signaling | Rabbit | #9701 | IHC | 1:500 |
| P19ARF | Abcam | Rabbit | ab80 | IF | 1:1000 |
| CK19 | Proteintech | Rabbit | 10712-1-AP | IF | 1:200 |

IB; immunoblot, IF; immunofluorescence, IHC; immunohistochemistry, IP; immunoprecipitation

**Supplementary Table 3. List of qRT-PCR primers used in this study**

| qRT-PCR primers | Sequence |
|---|---|
| RT_h45s rRNA Forward | ACCCACCCTCGGTGAGA |
| RT_h45s rRNA Reverse | CAAGGCACGCCTCTCAGAT |
| RT_h18s rRNA Forward | TTCGAACGTCTGCCCTATCAA |
| RT_h18s rRNA Reverse | ATGGTAGGCACGGCGACTA |
| RT_hGAPDH Forward | AAGAAGGTGGTGAAGCAGGC |
| RT_hGAPDH Reverse | TCCACCACCCTGTTGCTGTA |
| RT_hNAT10 Forward | ATAGCAGCCACAAACATTCGC |
| RT_hNAT10 Reverse | ACACACATGCCGAAGGTATTG |

**Supplementary Table 4. List of reagents used in this study**

| | | |
|---|---|---|
| Peroxidase AffiniPure Donkey Anti-Rabbit IgG (H+L) | Jackson ImmunoResearch | Cat# 711-035-152 |
| Peroxidase AffiniPure Donkey Anti-Mouse IgG (H+L) | Jackson ImmunoResearch | Cat# 715-035-150 |
| Agar | Lamda Biotech | Cat# C110 |
| RNaseOUT™ Recombinant Ribonuclease Inhibitor | Thermo Fisher | Cat# 10777019 |
| 14 mL, Sterile + Certified Free Open-Top Thinwall Polypropylene Tube, 14 x 95mm | Beckman Coulter | Cat# C14302 |
| Alcian Blue (pH 2.5) Stain Kit | Vector Laboratories | Cat# H-3501 |
| Corning™ BioCoat™ Collagen I, Rat Tail Culture Dish | Thermo Fisher | Cat# 08-774-8 |
| BSA | Sigma-Aldrich | Cat# A7906 |
| Pierce™ IP Lysis Buffer | Thermo Fisher | Cat# 87787 |
| Normal Rabbit IgG | Cell Signaling | Cat# 2729 |
| NuPAGE™ 4 to 12%, Bis-Tris, 1.5 mm, Mini Protein Gel | ThermoFisher | Cat# NP0335 |
| NuPAGE™ LDS Sample Buffer (4X) | ThermoFisher | Cat# NP0007 |
| SW 41 Ti Swining-Bucket Rotor | Beckman Coulter | Cat# 331362 |
| Beckman Optima LE-80K Ultracentrifuge | Beckman Coulter | Cat# 365668 |
| Histo-Clear | National Diagnostics | Cat# HS-200 |
| Primocin | InvivoGen | Cat# ant-pm-1,2 |
| trypLE™ Express | Gibco | Cat# 12605028 |
| PBS, 10X Sterile | Corning | Cat# 46-013-CM |
| Tricine | Sigma | Cat# T0377 |
| Triethanolamine | Sigma | Cat# 90278 |
| Cycloheximide | Sigma | Cat# C7698 |
| Sucrose | Sigma | Cat# S0389 |
| SSC Buffer 20× | Sigma | Cat# S6639 |
| Isoflurane | Covetrus | Cat# 11695067772 |
| Tris Buffered Saline, with Tween® 20, pH 8.0 | Sigma-Aldrich | Cat# P9039 |
| ProLong Gold antifade | Invitrogen | Cat# P36930 |

| | | |
|---|---|---|
| mountant with DAPI | | |
| SuperSignal™ West Pico PLUS Chemiluminescent Substrate | ThermoFisher | Cat# 34579 |
| Permount Mounting Medium | ThermoFisher | Cat# SP15-100 |
| PowerUp SYBR Green Master Mix | ThermoFisher | Cat# A25742 |
| RNase-Free DNase Set | Qiagen | Cat# 79254 |
| T-PER Tissue Protein Extraction Reagent | ThermoFisher | Cat# 78510 |
| Pierce™ BCA Protein Assay Kit | ThermoFisher | Cat# 23225 |
| NuPAGE™ LDS Sample Buffer (4X) | Invitrogen™ | Cat# NP0007 |
| 2-Mercaptoethanol | Sigma-Aldrich | Cat# M3148 |
| Pierce IP lysis buffer | ThermoFisher | Cat# 87787 |
| Dynabeads Protein A | ThermoFisher | Cat# 10001D |
| Hybond®-N+ Nylon Transfer Membrane | Amersham Life Sciences | |
| Northern blot starter kit | Roche | Cat# 12039672910 |
| CDP-Star® Chemiluminescent Substrate | Sigma | Cat# C0712 |
| PrimeScript™ RT Reagent Kit | Takara Bio Inc | Cat# RR037B |
| Triton X-100 | LabChem | Cat# LC262801 |
| Tamoxifen | Toronto Research Chemicals Inc | Cat# T00600 |
| Halt™ Protease and Phosphatase Inhibitor Cocktail, EDTA-free (100X) | ThermoFisher | Cat# 78443 |
| Donkey anti-Rabbit IgG (H+L) Highly Cross-Adsorbed Secondary Antibody, Alexa Fluor 594 | ThermoFisher | A-21207 |
| Donkey anti-Rabbit IgG (H+L) Highly Cross-Adsorbed Secondary Antibody, Alexa Fluor 647 | ThermoFisher | A-31573 |
| Donkey anti-Mouse IgG (H+L) Highly Cross-Adsorbed Secondary Antibody, Alexa Fluor 488 | ThermoFisher | A-21202 |

| | | |
|---|---|---|
| Donkey anti-Mouse IgG (H+L) Highly Cross-Adsorbed Secondary Antibody, Alexa Fluor 594 | ThermoFisher | A-21203 |
| Donkey anti-Mouse IgG (H+L) Highly Cross-Adsorbed Secondary Antibody, Alexa Fluor 647 | ThermoFisher | A-31571 |
| Donkey anti-Rabbit IgG (H+L) Secondary Antibody, Alexa Fluor 488, Invitrogen | ThermoFisher | A-21206 |
| Donkey anti-Goat IgG (H+L) Secondary Antibody, Alexa Fluor 594, Invitrogen | ThermoFisher | A-11058 |
| Donkey anti Rat IgG (H+L) Highly Cross-Adsorbed Secondary Antibody, Alexa Fluor 488 | ThermoFisher | A-21208 |
| Veriblot | Abcam | Cat# ab131366 |
| RPMI-1640 | Gibco | Cat# 11875093 |
| DMEM, high glucose | Gibco | Cat# 11965092 |
| Fetal Bovine Serum (FBS) | Gibco | Cat# 26140079 |
| RIPA Lysis and Extraction Buffer | ThermoFisher | Cat# 89900 |
| alamarBlue™ Cell Viability Reagent | Invitrogen™ | Cat# DAL1025 |
| Vectastain Elite ABC HRP Kit | Vector Laboratories | Cat# PK-6100 |
| DAB Substrate Kit | Thermo Scientific | Cat# 36000 |
| RNeasy Mini Kit | Qiagen | Cat# 74104 |
| Direct-zol RNA Miniprep Kits | Zymo Research Corporation | Cat # R2053 |
| Click-iT™ RNA Alexa Fluor™ 594 Imaging Kit | Invitrogen | Cat# C10330 |
| TRIzol™ Reagent | Thermo Fisher | Cat# 15596018 |
| Formaldehyde solution | Sigma-Aldrich | Cat# 252549 |
| Paraformaldehyde 16% Aqueous Solution EM Grade | Electron Microscopy Sciences | Cat# 15710 |
| AGS | ATCC | CRL-1739 |
| LS-174T | ATCC | CL-188 |
| Opti-MEM™ I Reduced Serum | Gibco | Cat# 31985070 |

| | | |
|---|---|---|
| Medium | | |
| Lipofectamine™ 2000 Transfection Reagent | Invitrogen™ | Cat# 11668019 |
| ChemiDoc™ MP Imaging System | Bio-Rad | Cat# 12003154 |
| Pannoramic MIDI II | Epredia | |
| AX R Confocal System with Eclipse Ti2-E Inverted Microscope | Nikon | |
| ImageJ | NIH | https://imagej.net/ |
| Prism 10 | GraphPad | https://www. graphpad.com/scientific-software/prism/ |
| Lionheart FX | BioTek, Agilent | |

**Supplementary Table 5. List of mouse strains used in this study**

| | | |
|---|---|---|
| B6.Cg-Tg(CAG-cre/Esr1*)5Amc/J | Jackson laboratories | RRID: IMSR_JAX: 004682 |
| B6.129-*Bhlha15*$^{tm3(cre/ERT2)Skz}$/J | Jackson laboratories | RRID: IMSR_JAX: 029228 |
| B6.129P2-*Trp53*$^{tm1Brn}$/J | Jackson laboratories | RRID: IMSR_JAX: 008462 |
| B6.129S2-*Trp53*$^{tm1Tyj}$/J | Jackson laboratories | RRID: IMSR_JAX: 002101 |
| B6.Cg-*Gt(ROSA)26Sor*$^{tm9(CAG-tdTomato)Hze}$/J | Jackson laboratories | RRID: IMSR_JAX:007909 |
| Nat10 flox/flox | Generated by the authors | |
| Nat10 heterozygous | Generated by the authors | |
